# Range expansion is both slower and more variable with rapid evolution across a spatial gradient in temperature

**DOI:** 10.1101/2023.09.14.557841

**Authors:** Takuji Usui, Amy L. Angert

## Abstract

Rapid evolution in colonizing populations can alter our ability to predict future range expansions. Recent theory suggests that the dynamics of replicate range expansions are less variable, and hence more predictable, with increased selection at the expanding range front. Here, we test whether selection from environmental gradients across space produces more consistent range expansion speeds, using the experimental evolution of replicate duckweed populations colonizing landscapes with and without a temperature gradient. We found that range expansion across a temperature gradient was slower on average, with range-front populations displaying higher population densities, and genetic signatures and trait changes consistent with directional selection. Despite this, we found that with a spatial gradient range expansion speed became more variable and less consistent among replicates over time. Our results therefore challenge current theory, highlighting that chance can still shape the genetic response to selection to influence our ability to predict range expansion speeds.

## INTRODUCTION

Predicting the movement of species ranges is a fundamental goal of ecology and is of increasing importance as biological invasions and climate change accelerate range expansion (Chen et al. 2011; Iseli et al. 2023; Nadeau & Urban 2019). Classic work on the ecological dynamics of range expansion has demonstrated that higher values of two key life-history traits – dispersal and reproductive output – increase the speed of range expansion (Kot et al. 1996; Skellam 1951). More recent work incorporating rapid evolution has shown that not only can dispersal and reproduction evolve to increase the average speed of range expansion, but that additionally, evolution can directly influence the amount of variability in the speed of population spread across replicate, range-expanding populations (Miller et al. 2020; Ochocki & Miller 2017; Williams et al. 2016; Wiliams et al. 2019). However, when and in what direction evolution modifies the variability of, and hence our ability to precisely predict, range expansions remain unclear.

On one hand, eco-evolutionary studies of range expansion predict that evolution should increase variability in range expansion speed across replicate populations. Specifically, serial founding events during range expansion can result in the fixation of maladaptive alleles at the range front and consequent reduction in fitness (‘expansion load’) through a spatial analogue of genetic drift (‘gene surfing’) (Excoffier et al. 2009; Klopfstein et al. 2006; Perrier et al. 2020). Gene surfing in small, range-expanding populations leads to the fixation of random amounts of expansion load across replicate populations, thus generating variability in range expansion speed (Miller et al. 2020; Phillips 2015; Williams et al. 2019). The implication then, is that any single realisation of range expansion in nature is drawn from a distribution of possible outcomes, making our ability to precisely predict the dynamics of range expansion more difficult.

In contrast, some recent experimental work demonstrates that evolution can also reduce variability in range expansion speed (Miller et al. 2020; Williams et al. 2019). This is predicted to occur when directional selection (e.g., for traits conferring greater dispersal or environmental tolerance; Chuang & Peterson 2016; Freedman et al. 2020; Lancaster et al. 2015) reduces trait variance at the range front, hence increasing consistency in the speed of range expansion (Phillips 2015; Williams et al. 2019). Overall, how evolution modifies the variability in range expansion speed may therefore depend on the relative strength of gene surfing and selection in range-front populations, and the factors that tip this balance one way or another (Miller et al. 2020; Urquhart-Cronish et al. 2022; Williams et al. 2019).

One key factor that could tip this balance between the variance-generating and variance-reducing effects of evolution is environmental variation in space. Specifically, species’ ranges in nature exist across broad-scale, spatial ecological gradients. Importantly, changing abiotic or biotic conditions across space could lead to slower range expansion due to the need for populations to adapt to the underlying spatial gradient. This initial maladaptation at the range front can allow more time for individuals to arrive from the range core, consequently increasing population size and genetic variation at the range front over time (Fronhofer et al. 2017; Gilbert et al. 2017). In these ‘steeper’ range boundaries, directional selection favouring traits that confer tolerance to stressful ecological conditions could offset any maladaptive gene flow from the range core, and importantly, weaken gene surfing and its genetic and demographic consequences (Gilbert et al. 2017; Williams et al. 2019). Thus, selection from underlying ecological gradients in space could make the speed of range expansion among replicate populations less variable and more predictable. Although there is a need to incorporate realistic sources of environmental variation, experimentally testing how selection from ecological gradients in space modifies the predictability of replicate range expansions in nature remains difficult (Miller et al. 2020).

Experimental range expansions in “miniaturised” landscapes (Larsen & Hargreaves 2020; Lustenhower et al. 2023) can provide a powerful bridge between theory and range expansion in natural landscapes. First, these landscapes differ from traditional microcosms in that replicate populations can disperse naturally across linear landscapes, such that realistic genetic and demographic spatial structure arises during range expansions. Second, through replicated landscapes we can track a distribution of ecological and evolutionary outcomes in colonising populations. This allows experimental insights into the factors that contribute to variability in range expansion that would not be possible without replication. Third, through experimental evolution we can track the eco-evolutionary dynamics of range expansion unfolding over multiple generations in real time, while field studies have mostly been retrospective, inferring past demographic and evolutionary processes based on current spatial patterns. Despite the increasing utility of these landscapes (Larsen & Hargreaves 2020), we still require experiments that test the dynamics of range expansion across landscapes that incorporate selection from spatial environmental variation.

Here, we use experimental populations of duckweeds in the *Lemna* species complex to provide a first experimental test for how selection from spatial abiotic gradients modifies variability in the rate of replicate range expansions. Using landscapes with and without a temperature gradient in space, we predicted that: (i) range expansion speed should be less variable among replicate populations moving across landscapes with a spatial temperature gradient than without. We predicted that this would occur as: (ii) range expansion across temperature gradients would be on average slower, resulting in higher population densities and steeper range boundaries. Consequently, we predicted that there should be stronger signatures of selection over genetic drift in populations expanding across temperature gradients. Specifically, we predicted: (iii) directional selection for traits relevant to temperature stress; and (iv) less among-replicate variability in genotypic composition than predicted by chance.

## MATERIALS & METHODS

### *Lemna* experimental populations

Duckweeds are small, freshwater angiosperms with a rapid generation time of 2–5 days under ideal conditions (Landolt 1986). Sexual reproduction is very rare in duckweeds, with reproduction at experimental timescales occurring exclusively through the clonal budding of daughter fronds (Docauer 1983). We sampled 20 accessions of duckweeds in the *Lemna* species complex from 20 sites around the Pacific Northwest region of North America (Table S1).

Microsatellite genotyping of each accession and molecular barcoding approach using Tubulin-Based Polymorphism (TBP) fingerprinting (Braglia et al. 2021a) detected that our accessions were composed of 11 unique genotypes, potentially grouping to 6 genotypes of *L. minor* and 5 genotypes of the putative *L. minor-L. turionifera* hybrid *L. japonica* (Appendix S1; Tables S1-S3). Species delineation remains challenging because these two lineages are ecologically and genetically similar to one another while varying widely in ploidy levels, and different molecular markers lead to inconsistent identification at the species level (Appendix S1). In our experiment, we therefore describe evolution as occurring through the sorting of unique genotypes within part of the *Lemna* species complex.

### Experimental landscapes and range expansion

We set up 50 experimental landscapes across five benches inside the UBC greenhouse, with each landscape constructed from aluminium gutters (150 cm x 12.5 cm; L x W; Fig. 1a). We set up a temperature treatment using submersible heaters (150W JBJ aquarium heaters) such that range expansion occurred across landscapes with a spatial gradient of increasing temperatures (“gradient landscapes” hereafter; β = 0.027, 95% CI: 0.027 to 0.028 °C/cm) or across landscapes with near-uniform temperatures (“uniform landscapes” hereafter; β = –0.001, 95% CI: –0.002 to 0.000 °C/cm; Fig. 1b). The position of gradient and uniform landscapes (*N* = 25 each) was randomised within each bench. We note that because we chose to standardize mean temperatures between our uniform and gradient landscapes (27.4°C; SD = 3.85°C and 26.9°C; SD = 1.53°C, respectively), temperatures at the start of gradient landscapes were relatively cooler than in uniform landscapes (24.3°C; SD = 2.6°C and 27.6°C; SD = 1.4°C, respectively; Fig. S1). However, we note that our outcomes still reflect the presence and absence of a temperature gradient across space in gradient and uniform landscapes, respectively (Appendix S2 for additional experimental details).

**Figure 1.**
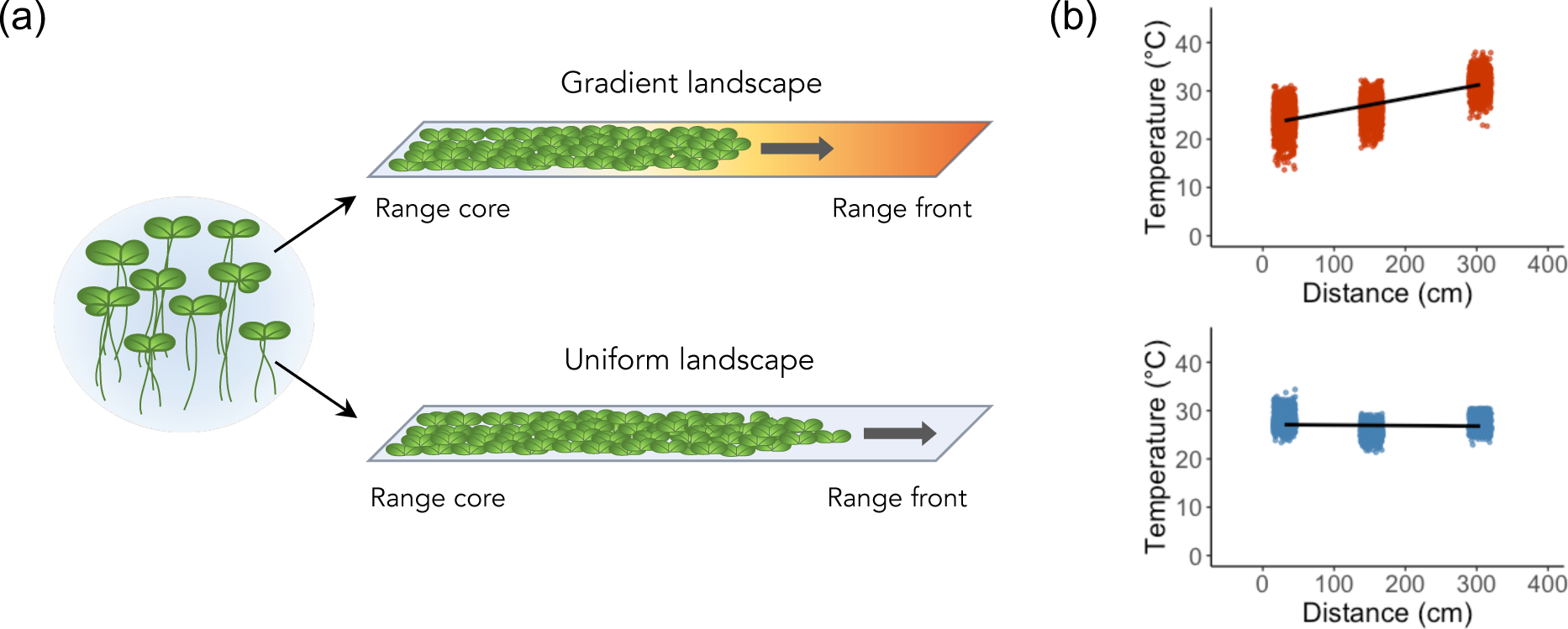
Experimental range expansion across landscapes with and without a spatial gradient in temperature. (a) Replicate, mixed-genotype populations of duckweed (*Lemna* species complex) expand their experimental ranges across increasingly warmer temperatures (blue to red) from the range core to the range front in gradient landscapes, or across benign and constant temperatures in uniform landscapes. (b) Observed temperatures across gradient (red) and uniform (blue) landscapes during the experiment. Lines represent the slopes of linear models of temperature by distance from range core. We found a significant and positive temperature slope in gradient landscapes (β_*+,-./’0_ = 0.027, 95% CI: 0.027 to 0.028 °C/cm; *t* = 126.49; *P* < 0.001) and a slightly negative slope in uniform landscapes (β_1’.23+4_ = –0.001, 95% CI: –0.002 to 0.000 °C/cm; t = 68.50; *P* < 0.01).

To initiate range expansion, we established genetically diverse *Lemna* populations, with each population consisting of an even density of 20 accessions (*N* = 20 individuals per accession for a founding population size of *N* = 400 individuals; Appendix S2). We seeded each population at one end of each landscape, allowing 2 days for plants to acclimate before range expansion. In gradient landscapes, we seeded populations at the cooler end of the landscape and allowed expansion into increasingly warmer temperatures (Fig. 1a). To allow all populations the opportunity to colonize the full spatial extent of the landscape, the experiment ran for 112 days. During our experiment, we lost one replicate due to structural damage and lost 10 replicates on one bench due to a disturbance event that disrupted population spread. We also lost 6 replicates due to heavy algal growth on the water surface that obstructed population spread. In total, we retained 33 experimental landscapes across four benches (*N* = 16 gradient landscapes; *N* = 17 uniform landscapes).

### Quantifying range expansion speed

We quantified the mean and variability in range expansion speed across landscapes for the initial 24 days (i.e., ∼5–12 generations) of range expansion. This timescale represents population spread during the exponential growth phase and thus is most consistent with theoretical models of range expansion (Kot et al. 1996; Skellam 1951). Also, some populations had reached the range limit of landscapes by day 24. These replicates would not be able to spread further and their inclusion after 24 days would artificially reduce the average and among-replicate variability in range expansion speed (i.e., the ceiling effect).

To quantify range expansion speed, we took aerial photographs of each landscape every 4 days (Nikkon D750). From these images, we first quantified the maximum distance travelled (cm) for each population by measuring the distance reached by the furthest individual every 4 days. We then quantified the mean and among-replicate variability in the distance travelled at each treatment level and time point. To estimate the difference in among-replicate variability of distance travelled for gradient and uniform landscapes, we calculated the log coefficient of variation ratio (lnCVR; Nakagawa et al. 2015) as:

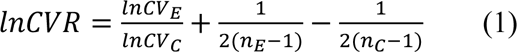

where subscripts *E* and *C* refer to the experimental (i.e., gradient) and control (i.e., uniform) landscapes, CV is the coefficient of variation, and *n* is the sample size. We calculated lnCVR such that positive values correspond to larger variability in distance travelled among gradient compared to uniform landscapes. We use lnCVR because CV accounts for the linear and positive mean-variance relationship on the log scale as observed in our dataset (Fig. S2) where shifts in variability could be driven by corresponding shifts in the mean (Taylor 1961). In such cases, it is preferable to use CV as a measure of variability to account for differences in the mean, although for sensitivity we also estimated the log variation ratio (lnVR) to estimate changes in variability (SD) without accounting for mean differences (Appendix S4). To report differences in CV, we back-transformed estimates of lnCVR.

### Quantifying range-front population density and steepness

We quantified spatiotemporal changes in population density at the range front by estimating the percent cover of live (i.e., green) fronds on the water surface. We did this by colour thresholding images obtained every 4 days using ImageJ (v1.53) and estimating percent cover in 2 cm x 12.5 cm grids at the range front (defined as the first 12 cm from the furthest travelled individual). We then fit a logistic model of percent cover across space for each landscape to quantify the maximum population density at the range front at each time point. We also quantified the steepness of the range boundary as the rate of increase in population density across space from the range front towards the range core for each landscape and time point. We did so by fitting an exponential model to the first three spatial estimates of percent cover closest to the range front of each landscape (i.e., percent cover at positions 0 cm, 2 cm, and 4 cm from the range front). Steepness was quantified as the exponent obtained from each model (Legault et al. 2020).

### Quantifying range-front evolution

We quantified evolution as the change in genotype frequencies in post-expansion populations compared to the founding population. Post-experiment, we haphazardly sampled 12 individual rafts of duckweeds from each replicate range front. Each sample was then genotyped using four microsatellite markers in a single multiplex reaction (Appendix S1; Table S2; Hart et al. 2017). We successfully obtained genotype sequences for 272 individuals (88% of all genotyped samples). To balance the number of samples obtained within each replicate and treatment, we rarefied samples and dropped replicates represented by less than *N* = 6 individuals due to DNA extraction or genotyping failure. For all downstream analyses, we therefore obtained a total of *N* = 6 individuals per replicate, for 10 and 17 replicates of gradient and uniform landscapes, respectively. We note a higher extraction and genotyping failure for samples obtained from gradient landscapes possibly due to greater damage to plant tissue from heat stress.

To investigate whether genotype sorting in the experiment led to trait evolution, we also measured a suite of genotype-specific traits *ex situ* in incubators and across a range of temperatures (Appendix S3; Fig. S5) including: maximum growth rate (*r*_max_); thermal optimum (*T*_opt_); critical thermal maximum (CT_max_); thermal performance breadth (i.e., the range of temperatures for which growth rate is within 80% of the maximum; Padfield et al. 2021); thermal tolerance breadth (i.e., the range of temperatures for which growth rate is positive); mean specific-leaf area (SLA; cm^2^/g); mean root-shoot ratio (cm/mg); plasticity in both SLA and root-shoot ratio (i.e., CV across temperature; Valladares et al. 2006); and mean raft number of duckweed fronds (i.e., the average number of fronds that stayed attached to each other during clonal reproduction to form multi-frond rafts). Mean raft number was used as a proxy for dispersal ability assuming plants with a smaller raft number would disperse further as fronds separate and release during budding. Before analyses, we scaled and centred each trait, and estimated mean trait values at the population level by weighting trait values by observed genotype frequencies.

### Statistical analyses

We tested for treatment differences in mean range expansion speed (prediction ii) by fitting a linear mixed-effects model using the *lme4* package (Bates et al. 2015). We fit maximum distance travelled (cm) as the response variable and an interaction between time (day) and temperature treatment (gradient *vs* uniform) as fixed predictors, where mean range expansion speed is modelled as slopes (cm/day). We also included bench ID as a fixed covariate (as there were only four bench levels; Gelman & Hill 2006) and fitted a random slope (i.e., speed) for each replicate. Next, we tested for treatment differences in among-replicate variability of distance travelled (prediction i) by fitting a linear model with lnCVR as the response variable and time as the predictor. We modelled the temporal trend in lnCVR as visual inspections of CV in distance travelled showed changes over time for both gradient and uniform landscapes (Fig. 2b). For sensitivity analyses, we also conducted a linear meta-regression of lnCVR to explicitly model sampling variation (Appendix S5). We then tested for treatment differences in range-front population density (i.e., carrying capacity) and steepness of the range boundary (i.e., exponent) (prediction ii) by fitting separate linear regressions with either population density or steepness as the response variable, respectively, and time, temperature treatment, and their interaction as predictors. For sensitivity analyses, we also conducted a separate meta-regression of population density and steepness by including the standard error around each observation using the package *metafor* (Viechtbauer 2010).

**Figure 2.**
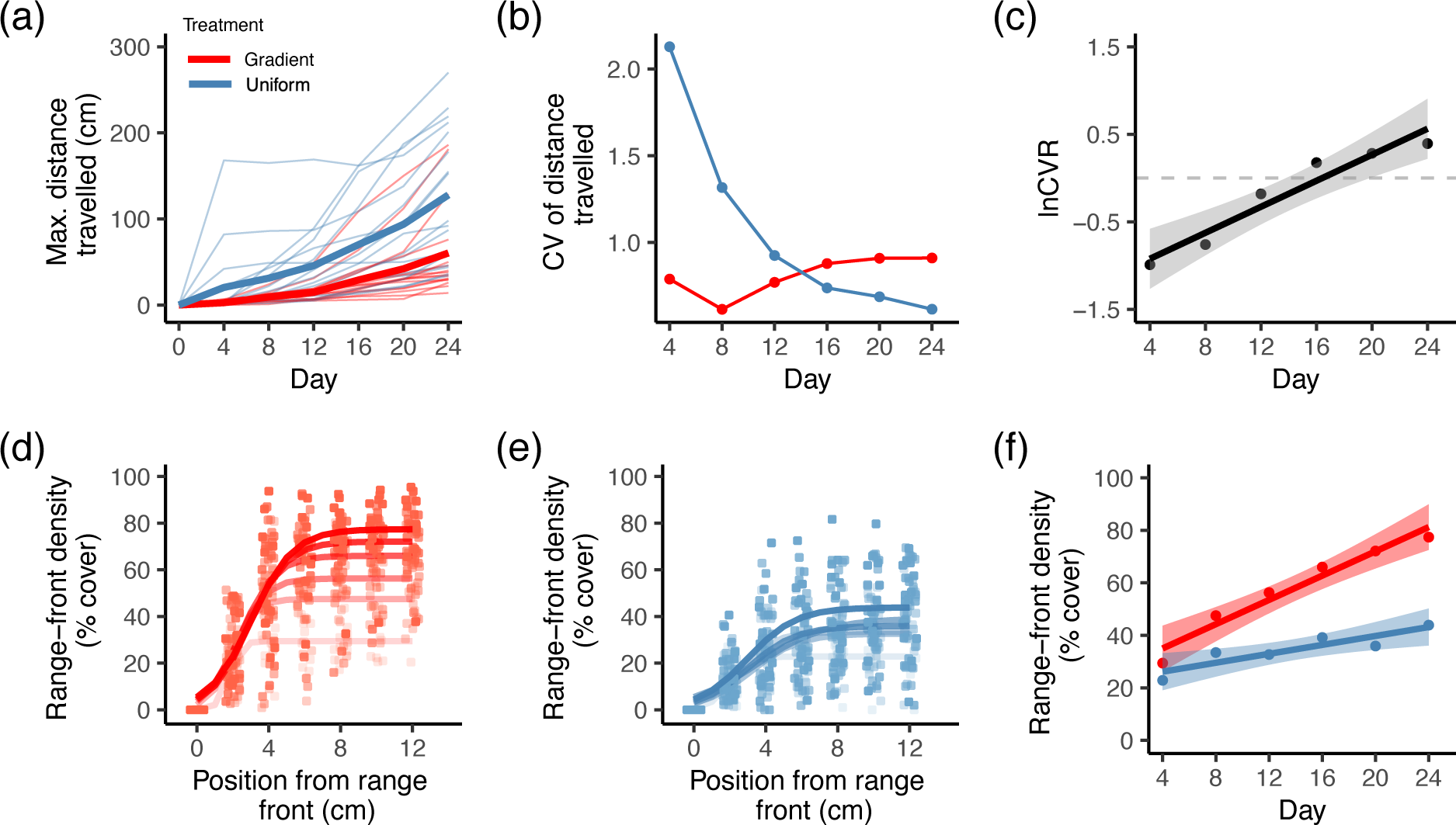
Population dynamics of range expansion across landscapes with (red) and without (blue) a temperature gradient across space. (a) Each line represents the maximum distance travelled over time for a single, replicate range expansion (mean values shown by thick lines). (b) Among-replicate variability (CV) in distance travelled over time for gradient and uniform landscapes. (c) Slope and 95% CI of changes in the log coefficient of variation ratio (lnCVR) over time, where estimates of lnCVR below zero (dashed grey line) indicate that there is less among-replicate variability (CV) across gradient than uniform landscapes. (d-e) Population densities at the range front (i.e., the first 0 to 12 cm) during range expansion. Lines show model predictions from a logistic regression of density across space, with increasing colour intensity representing changes over time from day 4 to 24. (f) Slope and 95% CI obtained from a linear regression of range-front density over time.

We next tested for treatment differences in the evolutionary dynamics of range expansion. First, we tested if evolution occurred during range expansion and if there were treatment differences in the amount of genotypic or trait change (prediction iii). We estimated genotypic or trait change in each landscape by quantifying the Euclidean distance between the initial and final genotype frequency or trait value in multidimensional genotype or trait space (i.e., across all 11 genotypes or 10 traits). We then fit separate linear regressions for genotype and trait change with Euclidean distance as the response variable, temperature treatment as the predictor, and bench ID as a fixed covariate. Upon observing evolution in multidimensional genotype frequencies and trait values, we conducted post-hoc tests to investigate univariate changes in specific genotypes and traits.

For genotypic change, we tested whether certain genotypes arose to higher frequencies than predicted by chance through a binomial test on the mean frequency of each genotype for each temperature treatment. For trait change, we estimated the mean and 95% CI of each trait for each temperature treatment and deemed traits to be significantly different from the founder population if the 95% CI of the final trait value did not overlap with the initial trait value.

Second, we also tested for treatment differences in the final genotype or trait composition in range-front populations (prediction iii). Here, we quantified compositional dissimilarity of genotypes or traits across all pairs of replicates within each treatment using Euclidean dissimilarity matrices, and then conducted separate PERMANOVAs with genotype or trait dissimilarity among replicates as the response variable, temperature treatment as the predictor, and bench ID as a fixed covariate. PERMANOVAs were conducted using the *adonis2* function in the *vegan* package (Okansen et al. 2022). To test for treatment differences in the mean values of each trait, we also conducted post-hoc *t*-tests on the final trait values observed in gradient and uniform landscapes.

Third, we tested if selection led to more consistent sets of genotypes at the range front and reduced among-replicate variability in genotype composition compared to a neutral model (prediction iv), or, if there was still a significant role of chance in shaping genotype composition at the range front. To do this, we compared the amount of observed among-replicate variability in each treatment to the simulated among-replicate variability predicted by random sampling. To quantify the amount of observed among-replicate variability, we averaged genotype frequencies across replicates within each treatment, calculated the Euclidean distance between this mean genotype composition and the observed genotype composition of each replicate (Fig. S7), and then summed these Euclidean distances across all replicates within each treatment. To prevent the sum value from being biased by differences in the number of replicates within each treatment, for this calculation only we randomly selected 10 uniform landscapes such that the number of replicates in uniform and gradient landscapes were equal (*N* = 10 for each). To quantify simulated among-replicate variability predicted by random sampling, we first pooled the genotype abundances across replicates within each temperature treatment (i.e., *N* = 60 individuals), randomly drew 6 individuals with replacement (i.e., representing the 6 genotypes in our observed data) and repeated this 10 times for each treatment (i.e., representing the 10 replicates of gradient and uniform landscapes). We then estimated the mean genotype composition for each temperature treatment, calculated the Euclidean distance between the simulated mean genotype composition and the simulated genotype composition of each replicate, and then finally summed these Euclidean distances for each treatment. We repeated this simulation a total of 100,000 times to estimate the probability of obtaining the observed among-replicate variability in genotype composition for each treatment based on random sampling.

Finally, we tested for treatment differences in the observed among-replicate variability through a permutation test, with Euclidian distances permuted randomly across treatment 100,000 times. All model-fitting was conducted in R (v4.3.0; R Core Team 2020).

## RESULTS

### Effects of temperature treatment on range expansion speed

On average, we found that range expansion speed across gradient landscapes was slower than across uniform landscapes (time by treatment effect: *P* = 0.016; Table S4; Fig. 2a), consistent with prediction (ii). Over 24 days of range expansion, the presence of a spatial temperature gradient reduced the average speed of range expansion by 50.9% (95% CI: 36.0% to 53.8%) compared to uniform landscapes (Table S4). We also found that the among-replicate variability in distance travelled differed between gradient and uniform landscapes (Fig. 2b and 2c). At the initial stages of range expansion, we found that there was less among-replicate variability (CV) in distance travelled across gradient compared to uniform landscapes (e.g., a 40.3% [95% CI: 28.6% to 56.8%] reduction in CV across gradient compared to uniform landscapes on day 4; Table S5), consistent with prediction (i). However, we found a significant increase in lnCVR over time (lnCVR slope: *P* = 0.002) such that by day 24, CV in distance travelled was 78.2% (95% CI: 28.6% to 151.0%) greater across gradient than uniform landscapes (Fig. 2c), contrary to prediction (i). This result was consistent in the sensitivity analysis including sampling variation around lnCVR in the linear model (Table S5). Sensitivity analysis of lnVR also showed that the slope of lnVR over time was positive (*P* = 0.002; Fig. S3) such that SD in distance travelled increased more in gradient compared to uniform landscapes over time.

### Effects of temperature treatment on range-front density and steepness

We found that the population density reached at the range front differed over time and between gradient and uniform landscapes (Fig. 2d-f). Although range-front densities were initially similar between gradient and uniform landscapes, population density increased more rapidly over time for gradient landscapes (treatment by time effect: *P* = 0.002; Fig. 2f), consistent with prediction (ii). By day 24, population density at the range front was 88.1% (95% CI: 76.5% to 104.0%) higher in gradient compared to uniform landscapes (Table S6). Along with increased population density, we also found range boundaries to be steeper in gradient landscapes (Table S7; Fig. S4), consistent with prediction (ii). These results were consistent in sensitivity models that included sampling error in estimates of range-front population density and steepness (Tables S6 and S7).

### Effects of temperature treatment on evolutionary change

We found that evolution in range-front populations occurred in both gradient and uniform landscapes, with significant changes in both genotype frequencies (Intercept: *P* < 0.001; Table S8; Fig. 3a) and trait values (Intercept: *P* = 0.002; Table S8; Fig. S6) in multidimensional genotype and trait space. We found support for prediction (iii) with genetic and trait signatures consistent with directional selection from a temperature gradient. First, range-front populations in gradient landscapes were often dominated by one genotype (G11; Fig. 3c). Post-hoc binomial tests showed that this genotype consistently increased in frequency in gradient landscapes (*P* < 0.001) but not in uniform landscapes (*P* = 0.869), suggesting selection for this genotype in the former (Fig. 3e). Second, final trait values for range-front populations in gradient landscapes showed significant changes in both thermal performance breadth and thermal tolerance breadth compared to the founder population (higher and lower values, respectively; Fig. 4; Table S10), although such a change was not observed in uniform landscapes. Third, we found that genotype composition of range-front populations differed between gradient and uniform landscapes after range expansion (PERMANOVA: *P* = 0.002; Table S9; Fig. 3b). Although differences in trait composition of range-front populations between gradient and uniform landscapes were only marginally significant (PERMANOVA: *P* = 0.066; Table S9; Fig. S6), post-hoc univariate *t*-tests on final trait values found reduced plasticity in root/shoot ratio in gradient compared to uniform range-front populations (*P* = 0.042; Table S11; Fig. 4). Importantly, despite these changes suggesting selection, and contrary to prediction (iv), we found that the observed among-replicate variability in genotype composition was greater than predicted by random sampling for both gradient (*P* < 0.001) and uniform (*P* < 0.001) landscapes (Fig. 5), suggesting an additional role of chance in shaping genotype composition at the range front.

**Figure 3.**
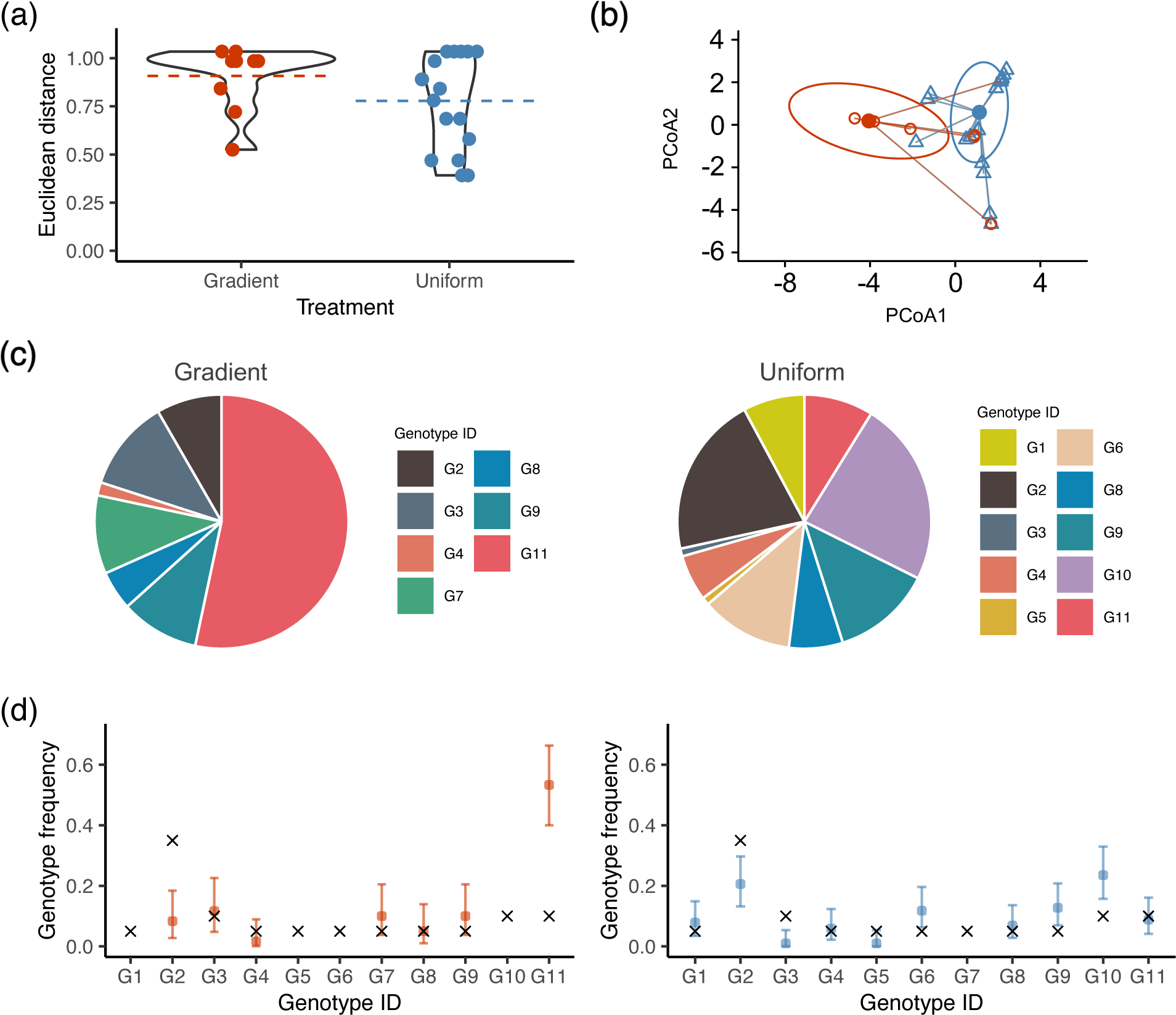
Evolutionary dynamics of range expansion across landscapes with and without a temperature gradient across space. (a) Violin plot for the magnitude of evolution observed in gradient (red) and uniform (blue) landscapes, as quantified by the Euclidean distance of genotype frequencies between founder and evolved populations at the range front. Points show the amount of change in genotype frequencies for each replicate, while dashed lines represent the mean change. (b) Genotype composition of range-front populations in gradient (red) and uniform (blue) landscapes. The first two axes of a Principal Coordinates Analysis (PCoA) are shown, accounting for 43.7% of the total variation in genotype composition. Filled circles are treatment centroids, while clear symbols are replicate populations in each treatment. Ellipses represent one standard deviation around treatment centroids, and lines represent distance from the centroid to each replicate population. (c) Mean genotype frequencies observed across range-front populations in gradient (left) and uniform (right) landscapes, with different colours representing unique genotypes. (d) Genotype frequency for gradient (left) and uniform (right) landscapes. Coloured points and error bars represent observed genotype frequencies and their 95% CIs in evolved range-front populations. The initial genotype frequency in the founder population is denoted by ‘x’.

**Figure 4.**
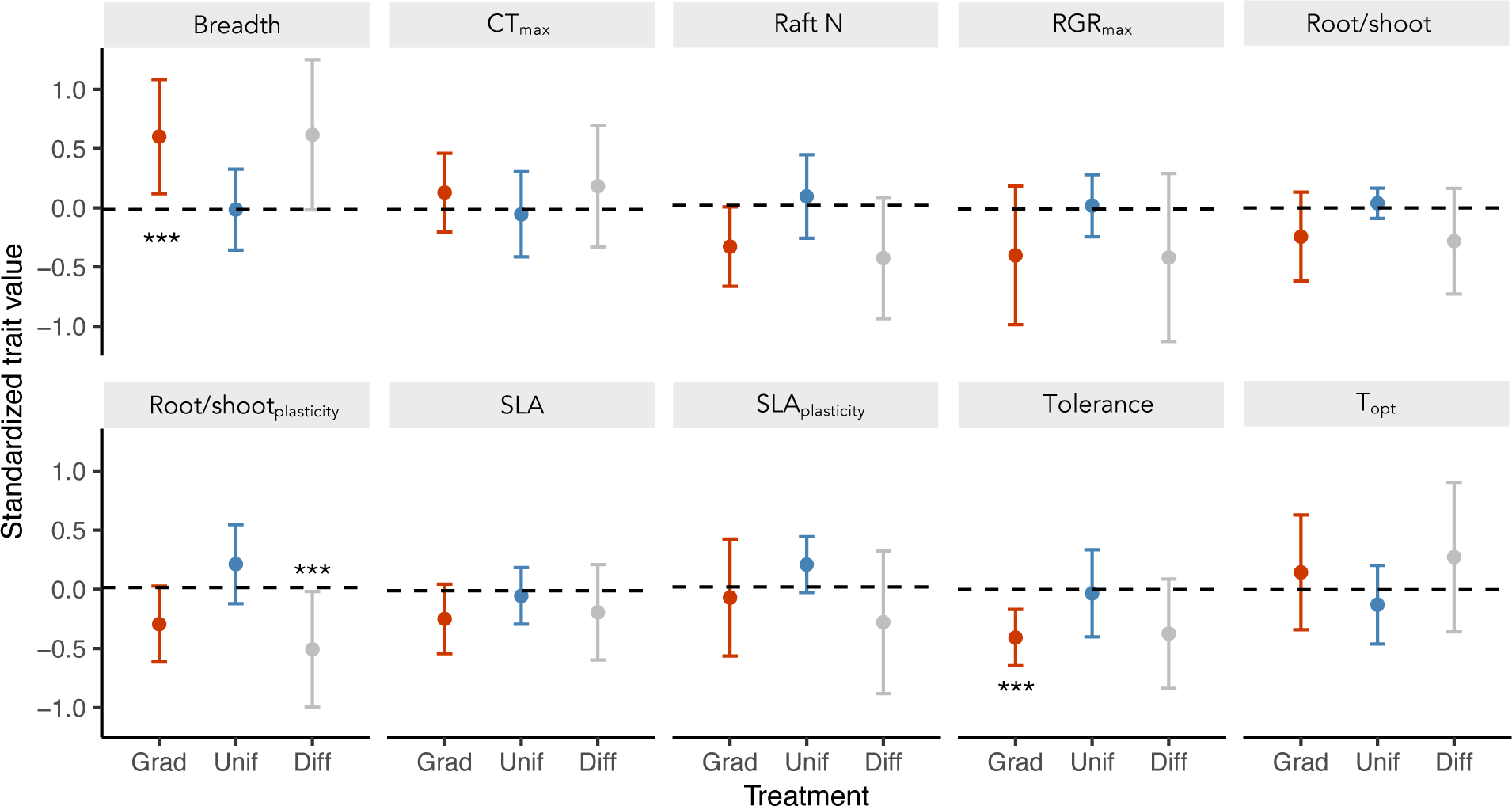
Evolved trait values for range-front populations in gradient and uniform landscapes. Red and blue points show mean trait values averaged across gradient and uniform replicates, respectively. The grey point shows mean treatment differences as estimated by gradient minus uniform trait values. Error bars show 95% confidence intervals. The horizontal dotted line represents the initial mean trait value of the founder population. *** Indicates traits that are significantly different from the founder population (if under red bars) or between gradient and uniform landscapes (if above grey bars). All trait values are weighted by genotype frequencies of each range-front population, and then scaled and centred.

**Figure 5.**
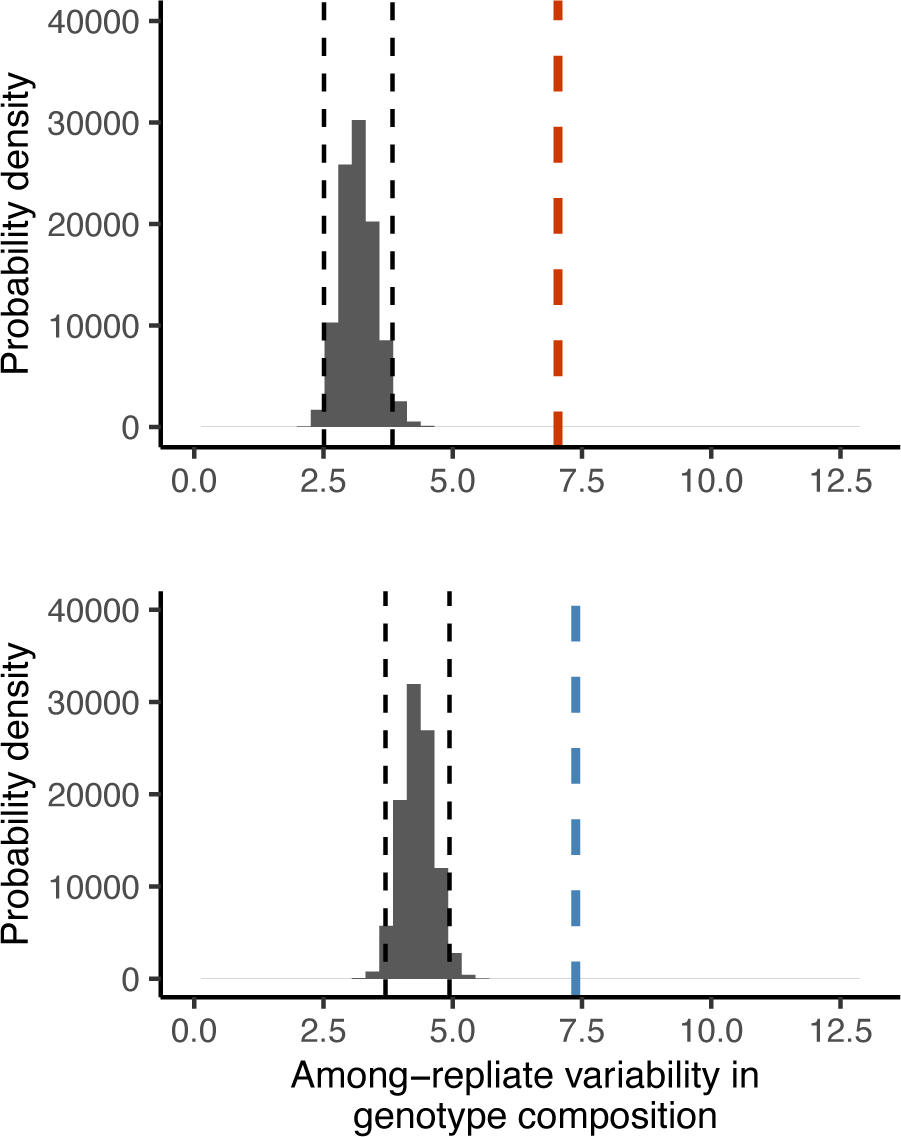
Observed versus simulated among-replicate variability in range-front genotype composition in gradient (top) and uniform (bottom) landscapes. Histograms represent the predicted probability distribution of among-replicate variability in genotype composition due to random sampling alone, with black dotted lines representing the 95% confidence interval. Vertical dashed red and blue lines show that the mean observed among-replicate variability in genotype composition for gradient and uniform landscapes are similar to each other (*P* = 0.741; Permutation test) but greater than predicted by random sampling (*P* < 0.001). Among-replicate variability in genotype composition was quantified as the summed Euclidean distance between the genotypic composition of each replicate population (*N* = 10 per treatment) and the mean genotypic composition of each treatment.

## DISCUSSION

Recent studies have highlighted that rapid evolution can complicate our ability to predict the eco-evolutionary dynamics of range expansions (Miller et al. 2020; Shaw et al. 2023; Williams et al. 2019; Zilio et al. 2023). Here, we used replicate range-expanding populations to empirically test whether the presence of selection from an abiotic gradient across landscapes could modify the among-replicate variability of, and hence our ability to precisely predict, range expansion speed. Notably, we found that over time range expansion speed became more variable among landscapes with a spatial temperature gradient than without (Fig. 2c). This occurred despite gradient landscapes having slower range expansions on average, greater range-front population densities (Fig. 2), and genetic and trait signatures consistent with directional selection (Fig. 3; Fig. 4) – conditions that would otherwise predict less gene surfing and thus more consistency in the speed of range expansions across replicate populations (Miller et al. 2020; Phillips 2015; Williams et al. 2019). Our results therefore challenge current theory and highlight the complexities of predicting range expansions even in highly standardised, replicate mesocosms with simple environmental variation.

We found that the greater variability in range expansion speed observed for gradient compared to uniform landscapes over time is driven by a combination of rapidly decreasing among-replicate variability (CV) in distance travelled for uniform landscapes, and slightly increasing CV for gradient landscapes over time (Fig. 2b). In uniform landscapes, the decrease in CV of distance travelled over time occurred as mean range expansion speed increased more quickly than the variance observed across replicates (Fig. 2a). A rapid increase in mean range expansion speed across uniform landscapes is consistent with previous experimental evolution work on range expansions, which found that range expansion speed can increase under constant environments due to spatial sorting for dispersal phenotypes and low-density selection for increased reproductive output at the range front (Ochocki & Miller 2017; Wagner et al. 2017; Weiss-Lehman et al. 2017). In contrast, for gradient landscapes, the slight increase in CV of distance travelled over time occurred as the variance in speed among replicates increased more quickly than the mean range expansion speed (Fig. 2a). This occurred as range expansion remained slow for most gradient landscapes but rapidly sped up in a few replicates (Fig. 2a), consequently elevating the among-replicate variability observed over time.

It is possible that the faster range expansion seen in some of these gradient landscapes was driven by selection from the temperature gradient (Moerman et al. 2022; Phillips et al. 2006; Szücs et al 2017; Williams et al. 2016). We found that gradient populations on average had higher frequencies of one genotype (G11) at the range front (Fig. 3c and 3d). In contrast, this genotype did not achieve frequencies >0.5 in any uniform landscapes (Fig. S8). We also found that gradient populations evolved higher thermal performance breadth (i.e., a broader range of temperatures across which plants can maintain near-optimal growth) but also lower thermal tolerance breadth (i.e., a narrower range of temperatures across which plants can exhibit positive growth; Fig. 4). The importance of maintaining performance across a broader breadth of temperatures has been long emphasised in studies of range expansion across latitudinal gradients in climate (Ackerly 2003; Lancaster 2016; Lancaster 2022). The concurrent evolution of lower thermal tolerance breadth is surprising and perhaps suggests a metabolic or evolutionary trade-off between maintaining high growth rates across a broader range of temperatures and the ability to withstand extreme temperature minimums or maximums, for example, if maintaining lower or upper thermal limits are costly (Barley et al. 2021). This also implies that studies on eco-evolutionary responses to temperature change need to consider evolution in the full shape of thermal performance curves to accurately predict outcomes of climate change (Malusare et al. 2022). Range-front populations in gradient landscapes also had significantly lower plasticity in root/shoot ratio compared to uniform landscapes indicating either a trade-off in responding to changes in temperature versus resource availability, or more consistent resource environments selecting for reduced plasticity, for example if there is constant competition for light and nutrients in these crowded and dense range fronts (Fig. 2d).

Despite the evidence above for selection in gradient landscapes, we also found that the among-replicate variability in genetic composition was greater than predicted by random sampling (Fig. 5). Such a pattern could result if chance still had a large role in shaping the genetic composition of range-expanding populations, for example, if individuals that initially occupy the range front through chance dispersal dominate subsequent range expansion (i.e., spatial priority effects; Hallatschek et al. 2007; Hewitt 2000). In combination with spatial environmental variation, spatial priority effects may shape the genotypes available to respond to selection in range-front populations, allowing non-optimal strategies to persist at the range front of some replicates (Baym et al. 2016; Williams et al. 2016). Indeed, further tests revealed that range expansion speed was significantly slower in gradient populations with no or low frequencies of the G11 genotype (slope = 1.832, 95% CI: 0.613 to 3.052 cm/day) compared to replicates with fixation or high frequencies of this genotype (slope = 4.516, 95% CI: 1.572 to 7.460 cm/day; Fig. S10). We therefore suggest that together with selection from a spatial environmental gradient, spatial priority effects and its variance-generating effects at the range front could lead to range expansion speeds that are slower on average, yet also increasingly different from one another over time. Definitively demonstrating the effects of spatial priority during range expansion would require future studies to track individual genotypes over a finer resolution in space and time (Baym et al. 2016).

Our study is, to our knowledge, the first to test how selection from a spatial environmental gradient alters the variability of replicate range expansions. We note that the dynamics observed in our clonal system are highly relevant to biological invasions given that clonal reproduction is common during range expansion and that many invasive plant species are capable of clonal spread (Cadotte et al. 2006; Song et al. 2013; Wang et al. 2017). Importantly, our study suggests that spatial priority effects, by blocking beneficial genotypes or mutants, could play a key role in governing the expansion dynamics of asexual organisms— especially so for populations undergoing unidirectional range expansions (e.g., spreading higher in elevation or latitude).

Although our study allows us to track a distribution of eco-evolutionary outcomes across highly replicated landscapes, we note that scaling up any experimental results from any micro-or mesocosm studies to natural populations requires careful consideration (Lustenhouwer et al. 2023). As populations traverse across environmentally variable and complex landscapes, studies of natural range expansions will need to incorporate how selection from underlying environmental variation in space can influence our ability to predict future range expansions.

## ACKNOWLEDGEMENTS

We thank H. Ahmed, M. Catalani, A. Doherty, K. Kubeck, E. Menchions, T. Scriber, and P. Seppala for their help with plant and greenhouse maintenance, and for providing assistance with experimental setup. We thank M. Chen, E. Lam, D. Moxley and A. Wong for their generous assistance with genetic work. We are grateful to R. Germain, S. Otto, M. Tseng, and J. Williams for their comments on earlier versions of the manuscript. Finally, we thank the three anonymous reviewers whose comments substantially improved this paper. This work was supported by the UBC International Doctoral Fellowship and the SSE Greg Lewontin Award to TU, and a Discovery Grant from the Natural Sciences and Engineering Research Council of Canada to ALA.

## Notes

### Competing Interest Statement

The authors have declared no competing interest.

### Summary of Updates

Manuscript revision after review.

## REFERENCES

Ackerly, D.D. (2003). Community assembly, niche conservatism, and adaptive evolution in changing environments. Int. J. Plant Sci., 164, S165–S184.

Appenroth, K.J., Teller, S. & Horn, M. (1996). Photophysiology of turion formation and germination in *Spirodela polyrhiza*. Biol. Plantarum, 38, 343–351.

Barley, J.M., Cheng, B.S., Sasaki, M., Gignoux-Wolfsohn, S., Hays, C.G., Putnam, A.B., et al. (2021). Limited plasticity in thermally tolerant ectotherm populations: evidence for a trade-off. Proc. R. Soc. B., 288: 20210765.

Bates, D., Mächler, M., Bolker, B. & Walker, S. (2015). Fitting linear mixed-effects models using lme4. J. Stat. Softw., 67, 1–48.

Baym, M., Lieberman T.D., Kelsic, E.D., Remy, C., Gross, R., Yelin, I. & Kishony, R. (2016). Spatiotemporal microbial evolution on antibiotic landscapes. Science, 353, 1147–1151.

Braglia, L., Breviario, D., Gianì, S., Gavazzi, F., De Gregori, J. & Morello, L. (2021a). New insights into interspecific hybridization in *Lemna* L. Sect. Lemna (Lemnaceae Martinov). Plants, 10, 2767.

Braglia, L., Lauria, M., Appenroth, K.J., Bog, M., Breviario, D., Grasso, A., et al. (2021b). Duckweed species genotyping and interspecific hybrid discovery by Tubulin-Based Polymorphism Fingerprinting. Front. Plant. Sci., 12, 625670.

Burton, O.J., Phillips, B.L. & Travis, J.M.J. (2010). Trade-offs and the evolution of life-histories during range expansion: Evolution during range expansion. Ecol. Lett., 13, 1210–1220.

Cadotte, M.W., Murray, B.R. & Lovett-Doust, J. (2006). Ecological patterns and biological invasions: Using regional species inventories in macroecology. Biol. Invasions, 8, 809–821.

Chen, I.C., Hill, J.K., Ohlemüller, R., Roy, D.B. & Thomas, C.D. (2011). Rapid range shifts of species associated with high levels of climate warming. Science, 333, 1024–1026.

Chuang, A. & Peterson, C.R. (2016). Expanding population edges: Theories, traits, and trade-offs. Glob. Change Biol., 22, 494–512.

Docauer, D. M. (1983). A nutrient basis for the distribution of the Lemnaceae. Dissertation. University of Michigan, Ann Arbor, Michigan, USA.

Excoffier, L., Foll, M. & Petit, R.J. (2009). Genetic consequences of range expansions. Annu. Rev. Ecol. Evol. S., 40, 481–501.

Freedman, M.G., Dingle, H., Strauss, S.Y., & Ramírez, S.R. (2020). Two centuries of monarch butterfly collections reveal contrasting effects of range expansion and migration loss on wing traits. Proc. Natl. Acad. Sci. U.S.A., 117, 28887–28893.

Fronhofer, E.A., Nitsche, N. & Altermatt, F. (2017). Information use shapes the dynamics of range expansions into environmental gradients. Glob. Ecol. Biogeogr., 26, 400–411.

Gelman, A. & Hill, J. (2006). Data analysis using regression and multilevel/hierarchical models. Cambridge University Press, Cambridge, UK.

Gilbert, K.J., Sharp, N.P., Angert, A.L., Conte, G.L., Draghi, J.A., Guillaume, F., et al. (2017). Local adaptation interacts with expansion load during range expansion: Maladaptation reduces expansion load. Am. Nat., 189, 368–380.

Hallatschek, O., Hersen, P., Ramanathan, S. & Nelson, D.R. (2007). Genetic drift at expanding frontiers promotes gene segregation. Proc. Natl., Acad. Sci. U.S.A., 104, 19926–19930.

Hart, S.P., Turcotte, M.M. & Levine, J.M. (2019). Effects of rapid evolution on species coexistence. Proc. Natl., Acad. Sci. U.S.A., 116, 2112–2117.

Healey, A., Furtado, A., Cooper, T. & Henry, R.J. (2014). Protocol: A simple method for extracting next-generation sequencing quality genomic DNA from recalcitrant plant species. Plant Methods, 27;10:21.

Hedges, L.V. & Nowell, A. (1995). Sex differences in mental test scores, variability, and numbers of high-scoring individuals. Science, 269, 41–45.

Hewitt, G. (2000). The genetic legacy of the Quaternary ice ages. Nature, 405, 907–913.

Huey, R.B. & Hertz, P.E. (1984). Is a Jack-of-All-Temperatures a Master of None? Evol., 38, 441–444.

Iseli, E., Chisholm, C., Lenoir, J., Haider, S., Seipel, T., Barros, A., et al. (2023). Rapid upwards spread of non-native plants in mountains across continents. Nat, Ecol. Evol., 7, 405–413.

Klopfstein, S., Currat, M. & Excoffier, L. (2006). The fate of mutations surfing on the wave of a range expansion. Mol. Biol. Evol., 23, 482–490.

Kot, M., Lewis, M.A. & Van Den Driessche, P. (1996). Dispersal data and the spread of invading organisms. Ecol., 77, 2027–2042.

Lancaster, L.T., Dudaniec, R.Y., Hansson, B. & Svensson, E.I. (2015). Latitudinal shift in thermal niche breadth results from thermal release during a climate-mediated range expansion. J. Biogeogr., 42, 1953–1963.

Lancaster, L.T. (2016). Widespread range expansions shape latitudinal variation in insect thermal limits. Nat. Clim. Change, 6, 618–621.

Lancaster, L.T. (2022). On the macroecological significance of eco-evolutionary dynamics: The range shift–niche breadth hypothesis. Philos. T. R. Soc. B., 377, 20210013.

Landolt, E. (1986). Biosystematic investigations in the family of duckweeds, Lemnaceae: the family of Lemnaceae, a monographic study. Volume 2: Morphology, karyology, ecology, geographic distribution, systematic position, nomenclature descriptions. Veroeffentlichungen des Geobotanischen Institutes der ETH, Stiftung Rubel, Zurich, Switzerland.

Larsen, C.D. & Hargreaves, A.L. (2020). Miniaturizing landscapes to understand species distributions. Ecography, 43, 1625–1638.

Legault, G., Bitters, M.E., Hastings, A. & Melbourne, B.A. (2020). Interspecific competition slows range expansion and shapes range boundaries. Proc. Natl. Acad. Sci. U.S.A., 117, 26854–26860.

Lustenhouwer, N., Moerman, F., Altermatt, F., Bassar, R.D., Bocedi, G., Bonte, D., et al. (2023). Experimental evolution of dispersal: Unifying theory, experiments and natural systems. J. Anim. Ecol., 92, 1113–1123.

Malusare, S.P., Zillio, G. & Fronhofer, E.A. (2022). Evolution of thermal performance curves: A meta-analysis of selection experiments. J. Evol. Biol., 36, 15–28.

Miller, T.E.X., Angert, A.L., Brown, C.D., Lee-Yaw, J.A., Lewis, M., Lutscher, F., et al. (2020). Eco-evolutionary dynamics of range expansion. Ecol., 101. e03139.

Moerman, F., Fronhofer, E.A., Altermatt, F. & Wagner, A. (2022). Selection on growth rate and local adaptation drive genomic adaptation during experimental range expansions in the protist *Tetrahymena thermophila*. J. Anim. Ecol., 91, 1088–1103.

Nadeau, C.P. & Urban, M.C. (2019). Eco-evolution on the edge during climate change. Ecography, ecog.04404.

Nakagawa, S., Poulin, R., Mengersen, K., Reinhold, K., Engqvist, L., Lagisz, M., et al. (2015). Meta-analysis of variation: Ecological and evolutionary applications and beyond. Methods Ecol. Evol., 6, 143–152.

Ochocki, B.M. & Miller, T.E.X. (2017). Rapid evolution of dispersal ability makes biological invasions faster and more variable. Nat. Commun., 8, 14315.

Oksanen, J., Simpson, G.L., Blanchet, F.G., Kindt, R., Legendre, P. & Minchin, P.R. (2022). Vegan: Community ecology package (v 2.6–2).

Pachepsky, E. & Levine, J.M. (2011). Density dependence slows invader spread in fragmented landscapes. Am. Nat., 177, 18–28.

Padfield, D., O’Sullivan, H. & Pawar, S. (2021). rTPC and nls.multstart: A new pipeline to fit thermal performance curves in R. Methods Ecol. Evol., 12, 1138–1143.

Perrier, A., Sánchez-Castro, D. & Willi, Y. (2020). Expressed mutational load increases toward the edge of a species’ geographic range. Evol., 74, 1711–1723.

Phillips, B.L. (2015). Evolutionary processes make invasion speed difficult to predict. Biol. Invasions, 17, 1949–1960.

Phillips, B.L., Brown, G.P., Webb, J.K. & Shine, R. (2006). Invasion and the evolution of speed in toads. Nature, 439, 803–803.

R Core Team (2020). R: A language and environment for statistical computing. R Foundation for Statistical Computing, Vienna, Austria.

Senevirathna, K.M., Crisfield, V.E., Burg, T. & Laird, R.A. (2021). Hide and seek: Molecular barcoding clarifies the distribution of two cryptic duckweed species across Alberta. Botany, 99, 795–801.

Shaw, A.K., Lutscher, F. & Popovic, L. (2023). Pushed to the edge: Spatial sorting can slow down invasions. Ecol. Lett., 26, 1293–1300.

Skellam, J.G. (1951). Random Dispersal in Theoretical Populations. Biometrika, 38, 196–218.

Slatyer, R.A., Hirst, M. & Sexton, J.P. (2013). Niche breadth predicts geographical range size: A general ecological pattern. Ecol. Lett., 16, 1104–1114.

Song, Y.B., Yu, F.H., Keser. L.H., Dawson, W., Fischer, M., Dong, M. & van Kleunen, M. (2013). United we stand, divided we fall: A meta-analysis of experiments on clonal integration and its relationship to invasiveness. Oecologia, 171, 317–327.

Szűcs, M., Vahsen, M.L., Melbourne, B.A., Hoover, C., Weiss-Lehman, C. & Hufbauer, R.A. (2017). Rapid adaptive evolution in novel environments acts as an architect of population range expansion. Proc. Natl. Acad. Sci. U.S.A., 114, 13501–13506.

Urquhart-Cronish, M., Angert, A.L., Otto, S.P. & MacPherson, A. (2022). Density-dependent selection during range expansion affects expansion load in life-history traits. bioRxiv, 2022.11.08.515702.

Valladares, F., Sanchez-Gomez, D. & Zavala, M.A. (2006). Quantitative estimation of phenotypic plasticity: bridging the gap between the evolutionary concept and its ecological applications. J. Ecol., 94, 1103–1116.

Viechtbauer, W. (2010). Conducting meta-analyses in R with the metafor package. J. Stat. Softw., 3, 1–48.

Volkova, P.A., Nachatoi, V.A. & Bobrov, A.A. (2023). Hybrid between *Lemna minor* and *L. turionifera* (*L. × japonica*, Lemnaceae) in East Europe is more frequent than parental species and poorly distinguishable from them. Aquat. Bot., 184, 103593.

Wagner, N.K., Ochocki, B.M., Crawford, K.M., Compagnoni, A. & Miller, T.E.X. (2017). Genetic mixture of multiple source populations accelerates invasive range expansion. J. Anim. Ecol., 86, 21–34.

Wang, Y.J., Müller-Schärer, H., van Kleunen, M., Cai, A.M., Zhang, P., Yan, R., et al. (2017). Invasive alien plants benefit more from clonal integration in heterogeneous environments than native. New Phytol., 216, 1072–1078.

Weiss-Lehman, C., Hufbauer, R.A. & Melbourne, B.A. (2017). Rapid trait evolution drives increased speed and variance in experimental range expansions. Nat. Commun., 8, 14303.

Williams, J.L., Kendall, B.E. & Levine, J.M. (2016). Rapid evolution accelerates plant population spread in fragmented experimental landscapes. Science, 353, 482–485.

Williams, J.L., Hufbauer, R.A. & Miller, T.E.X. (2019). How evolution modifies the variability of range expansion. Trends Ecol. Evol., 34, 903–913.

Zilio, G., Krenek, S., Gougat-Barbera, C., Fronhofer, E.A. & Kaltz, O. (2023). Predicting evolution in experimental range expansions of an aquatic model system. Evol. Lett., 7, 121–131.

