## Supplementary Information for "Range expansion is both slower and more variable with rapid evolution across a spatial gradient in temperature"

**Supplementary information contents:**

Supplementary Appendix
- Appendix S1. DNA extraction, genotyping, and species delimitation.
- Appendix S2. Additional experimental details.
- Appendix S3. Quantifying genotype-specific traits.
- Appendix S4. Sensitivity analyses of lnVR over time.
- Appendix S5. Sensitivity analyses of sampling variation in lnCVR.

Supplementary Figures
- Figure S1. Mean temperatures at the starting locality of the experiment.
- Figure S2. Linear relationship between mean and variability in distance travelled.
- Figure S3. Slope and 95% CI for change in lnVR of expansion speed over time.
- Figure S4. Steepness of range boundary in gradient and uniform landscapes.
- Figure S5. PCA of trait space and thermal performance curves across genotypes.
- Figure S6. Trait change and composition in gradient and uniform landscapes.
- Figure S7. Euclidean distances to the mean genotype frequency of each treatment.
- Figure S8. Within-replicate genotype composition in gradient and uniform landscapes.
- Figure S9. Composition of *L. minor* and *L. japonica* at the range front.
- Figure S10. Effect of G11 on mean speed in gradient landscapes.

Supplementary Tables
- Table S1. Sampling locations and coordinates of *Lemna* accessions.
- Table S2. Microsatellite markers used for multiplex genotyping of accessions.
- Table S3. Primer sequences used for barcoding accessions to species-level.
- Table S4. Model coefficients for mean range expansion speed.
- Table S5. Model coefficients for among-replicate variability in range expansion speed.
- Table S6. Model coefficients for range-front population density.
- Table S7. Model coefficients for range-front steepness.

Table S8. Model coefficients for magnitude of genotypic and trait change.
Table S9. PERMANOVA model coefficients for genotype and trait composition.
Table S10. Evolved trait values in uniform and gradient landscapes.
Table S11. Model coefficients from *t*-tests on treatment differences in trait means.
Table S12. Model coefficients for magnitude of genotype change at range core and edge.

**Appendix S1: DNA extraction, genotyping and species delimitation**

To obtain DNA from our duckweed samples, we used a modified CTAB protocol (Healey et al. 2014). Briefly, we first flash-froze and then lysed plant tissue from each sample by shaking samples vigorously with 1.3 mm chrome-steel beads (Biospec Products, Bartlesville, OK), CTAB buffer and 0.5% β-mercaptoethanol. Next, we separated out the aqueous layer from the organic layer by centrifuging with chloroform-isoamyl alcohol, and subsequently precipitated out the DNA by centrifuging with isopropanol. Lastly, DNA pellets were washed with 70% ethanol, and then resuspended with 10 mM TrisHCl before quantification, genotyping and barcoding.

To genotype each sample, we used four unique and fluorescently labelled microsatellite markers developed by Hart et al. (2019) (Table S2). We used a 15 µL multiplex PCR, which included 1 unit of 5X Taq Master Mix (New England BioLabs, Ipswich, MA), 0.5 µL of DNA template, and between 0.14 µM and 0.26 µM of each primer pair. We ran a touchdown PCR with an initial denaturation at 95°C for 2 mins, followed by 5 cycles of denaturation (95°C, 30 secs), annealing (60 °C, 1 min; decreasing 1°C per cycle), and extension (68°C, 30 secs). We followed this with 30 cycles with an annealing temperature of 55°C and a final extension at 68°C for 5 mins. Post-cycling, we conducted fragment length analyses on an ABI 3730XL genetic analyzer (Thermo Fisher Scientific, Waltham, MA) at the UBC Sequencing and Bioinformatics Consortium. Finally, we visualised, cleaned, and fit ladders and loci on each sample using Geneious Prime (v 2023.1.0).

To barcode each sample, experimental plant accessions were sent to the Rutgers Duckweed Stock Cooperative for Tubulin-Based Polymorphism (TBP) fingerprinting. We used TBP markers as recent findings show that species barcodes based on maternally inherited plastid loci may have difficulty distinguishing the cryptic *Lemna* species complex, particularly between *L. minor* and *L. japonica* (Braglia et al. 2021b). We used primers designed by Braglia et al. (2021a) targeting the $\beta$-tubulin gene (Table S3) with PCR conditions as follows: initial denaturation at 95°C for 3 minutes, followed by 35 cycles of denaturation (95°C, 50 secs), annealing (52°C, 50 secs), and extension (72°C, 1 min), and then a final extension at 72°C for 3 minutes. Post-cycling, products were visualized on 1.2% agarose gels (100V at 45 minutes).

Duckweeds in the *Lemna* species complex include *L. minor*, *L. turionifera*, and *L. japonica*, with some recent studies proposing *L. japonica* to be a hybrid between *L. minor* and *L. turionifera* (Braglia et al. 2021a; Braglia et al. 2021b; Volkova et al. 2023). These lineages are ecologically and genetically similar to one another and can vary widely in ploidy levels, making species delineation challenging. Different molecular markers can also lead to inconsistent identification at the species level, with for example, our initial molecular barcoding based on new plastid markers (Senevirathna et al. 2021) showing that our experimental genotypes were a mix of *L. minor* and *L. turionifera*. Nonetheless, analysis of our genetic data at the "species" level shows that range-front populations in gradient landscapes have a higher mean frequency of the putative hybrid *L. japonica* and a lower mean frequency of *L. minor*, compared to both the range-front population in uniform landscapes and the founder population (Fig. S9). However, this seems to be driven by an increase in a particular genotype (G11) rather than being through a general increase of all *L. japonica* genotypes (Fig. S9). We note that changes at the genotype level (i.e., evolution) therefore occurred in spite of any potential changes at higher-level groupings.

**Appendix 2: Additional experimental details**

Upon sampling duckweed plants from natural ponds, we immersed fronds in a dilute bleach solution (10% v:v) to create axenic plants of each accession. We maintained each of our 20 accessions in 250 mL erlenmeyer flasks with 100 mL of artificial pond media (Appenroth et al. 1996), inside a temperature-controlled laboratory (20°C) and under full-spectrum LED lighting (SunBlaster 6400K; 16:8 hour light:dark cycle). To obtain sufficient numbers of individuals for the experimental founder population, we grew out each accession for 4 weeks inside 20 L buckets at the UBC Botany greenhouse, with 17 L of water, 4 L of autoclaved potting soil (MiracleGro Potting Mix), 1.32 mM of $KNO_3$, 0.16 mM of $Ca(NO_3)_2$, and 0.02 mM of $KH_2PO_4$ as fertilisers.

In our experimental landscapes, each landscape was filled with 13.5 L of water and 2.5 L of potting soil. To ensure nutrients and water-levels did not deplete over time, we added 0.13 mM of $KNO_3$, 0.02 mM of $Ca(NO_3)_2$, and 0.002 mM of $KH_2PO_4$ every 4 days to each landscape, and continuously regulated water-levels using a float-valve connected to a sump pump. We connected drip emitters to the sump pump so that water refilling occurred slowly and did not disrupt the spatial structure of populations. We monitored water temperatures using three underwater temperature loggers (HOBO pendants) placed at equal spacing within each landscape. To establish a temperature gradient, we used a submersible heater (150W JBJ aquarium heaters) set to 37°C and placed at one end of each gradient landscape. To maintain uniform temperatures, we placed two heaters set to 24°C at both ends of each uniform landscape. These set temperatures were chosen as they are close to the critical thermal maxima ($T_{max}$) and thermal optima ($T_{opt}$), respectively, of the *Lemna* genotypes used (Fig. S5).

We note that because we chose to standardize mean temperatures between our uniform and gradient temperatures, temperature differences at the initial stages of range expansion (i.e., when response to selection could be strongest due to the available standing genetic diversity in the founder population) could potentially drive differences in range expansion dynamics. Even so, comparison to a subset of genetic data obtained post-range expansion at the range core show no significant changes in genotype frequency from the founder population (Table S12) suggesting that selection due to temperature differences at the range core was minimal and that the temperature gradient experienced during range expansion drove evolutionary dynamics.

### Appendix S3. Quantifying genotype-specific traits

We quantified genetic variation in 10 traits using three incubators programmed at a range of temperatures (Panasonic MIR 254 PA Cooled Incubators with LED lighting set at 16:8 hour light:dark cycle). In each incubator, we first measured growth rates of each genotype across eight temperature levels (5, 10, 15, 20, 25, 30, 35, and 40 °C), randomising the temporal sequence of set temperatures over time. For each temperature level, we first seeded ~10 (SD = 3.5) individuals of each genotype inside clear plastic cups containing 100 mL of artificial pond media (Appenroth et al. 1996). We placed $N = 2$ cups per genotype inside each incubator, and therefore had a total of $N = 6$ replicates per genotype per temperature level. We counted the number of live individuals on days 0 and 8 to estimate the per capita growth rate ($r_{ij}$) for each genotype $i$ at each temperature $j$ using the equation:

$$r_{ij} = \frac{\log(N_{t2}) - \log(N_{t1})}{t_2 - t_1}$$

where $N_{t2}$ and $N_{t1}$ are the number of individuals on days 8 and 0, respectively. To fit thermal performance curves and estimate the thermal optima and maxima for each of our 11 genotypes, we used nonlinear least squares (NLS) regression to fit five common thermal performance functions using the package *rTPC* (Padfield et al. 2021). For each genotype, we then picked the best fit model using AIC scores and derived five genotype-specific parameters for analysis: (i) temperature optimum ($T_{opt}$); (ii) thermal performance breadth; (iii) thermal tolerance breadth; (iv) critical thermal maximum ($CT_{max}$); and (v) maximum growth rate ($RGR_{max}$).

Additionally, for each genotype and temperature level, we estimated the mean root length (cm) by sampling the longest root of a haphazardly chosen plant within each cup. We estimated the mean frond area ($cm^2$) by quantifying the total area of live fronds within each cup and dividing by the total number of individuals. We also estimated mean frond dry mass (mg) by harvesting roots of all individuals from each cup, drying in a 70 °C oven for at least 24 hours, and dividing by the total number of individuals. From these measurements, we then quantified two composite traits for each genotype for analysis: specific-leaf area (SLA) as the ratio of frond area to frond dry mass ($cm^2/g$) and the ratio of root length to frond dry mass (cm/mg; hereafter “root/shoot” ratio). We also estimated plasticity in SLA and root/shoot ratio by calculating the coefficient of variation (CV) in each trait across temperatures (Valladares et al. 2006). Finally, we quantified the tendency of duckweeds to stay attached to form multi-frond rafts during reproduction versus its tendency to release fronds by breaking the stipule. We measured this for each genotype by estimating the mean number of fronds (mean raft number) in a single raft across benign (20-25°C) and hot temperatures (35-40°C). Mean raft number was used as a proxy for dispersal ability assuming that plants with smaller mean raft number would disperse further due to the tendency to separate and release of fronds.

**Appendix S4. Analysis of lnVR over time**

In addition to lnCVR, we estimated changes in the log variation ratio (lnVR) over time, which is the ratio of standard deviations (SD) between gradient and uniform groups (Hedges & Nowell 1995). We estimated lnVR as:

$$lnVR = \frac{lnSD_E}{lnSD_C} + \frac{1}{2(n_E-1)} - \frac{1}{2(n_C-1)}$$

where subscripts *E* and *C* refer to the experimental (i.e., gradient) and control (i.e., uniform) landscapes, and *n* is the sample size. We calculated lnVR such that positive values correspond to larger SD in distance travelled among gradient compared to uniform landscapes. We note that changes in SD and therefore lnVR can be driven by changes in the mean, where greater means can correspond to greater SDs as found in our dataset (Fig. S2). Despite this, consistent to analyses of lnCVR, we also find a positive slope in lnVR over time, which translates to SD increasing faster over time in gradient compared to uniform landscapes (Fig. S3).

**Appendix S5. Sampling variation in lnCVR and sensitivity analysis**

We estimated the sampling variance around lnCVR ($s^2_{\mathrm{lnCVR}}$) using equation 12 in Nakagawa et al. (2015):

$$s^2_{lnCVR} = \frac{s_C^2}{n_c \bar{x}_c^2} + \frac{1}{2(n_c-1)} - 2p\sqrt{\frac{s_C^2}{n_c \bar{x}_c^2} * \frac{1}{2(n_c-1)}}$$

$$+ \frac{s_E^2}{n_E \bar{x}_E^2} + \frac{1}{2(n_E-1)} - 2p\sqrt{\frac{s_E^2}{n_E \bar{x}_E^2} * \frac{1}{2(n_E-1)}}$$

where subscripts *E* and *C* refer to the experimental (i.e., gradient) and control (i.e., uniform) groups, CV corresponds to the coefficient of variation, *n* is the sample size, *s* is the standard deviation (SD), *x*-bar is the mean, and *p* is the correlation between the mean and SD on the log scale (Nakagawa et al. 2015). For sensitivity analyses of changes in lnCVR over time, we included the sampling variance around lnCVR in a meta-regression using the *metafor* package (Viechtbauer et al. 2010), with time (day) as a fixed predictor and effect size ID as a random effect.

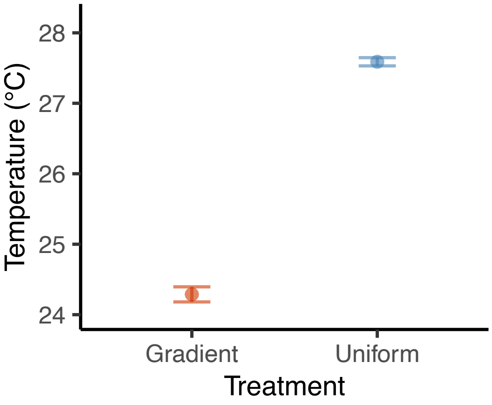


**Figure S1:** Differences in temperatures at the starting locality of the experiment. Points are the mean temperatures, while error bars represent the 95% CI. Note that the y-axis is truncated and does not start at zero.

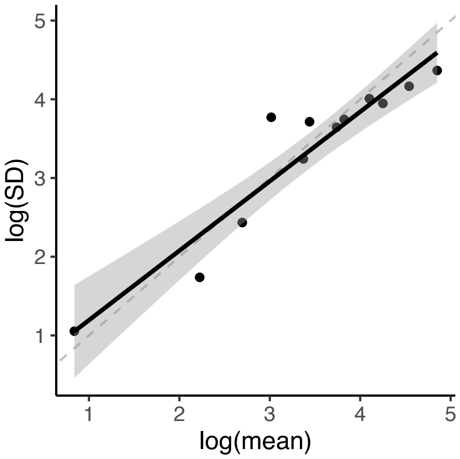


**Figure S2:** Relationship between mean and variance on the log scale (i.e., Taylor's Law). Data points represent the mean and SD in the maximum distance travelled for all landscapes, between day 4 and 24 (estimates obtained every 4 days). The slope and 95% CI from a linear regression is plotted ($\beta = 0.883$, 95% CI: 0.670 to 1.095), with the 1:1 relationship shown by the dashed line.

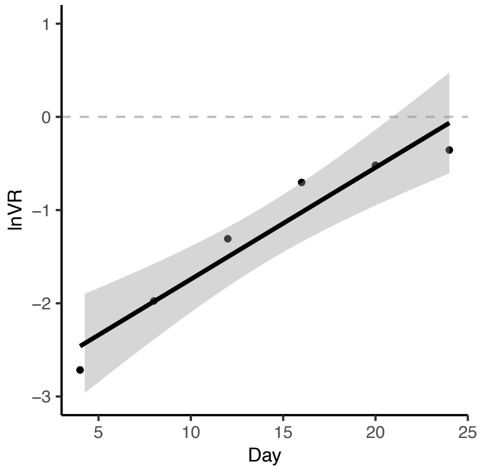


**Figure S3:** Slope and 95% CI from a linear model estimating the change in log variation ratio (lnVR) over time ($\beta$ = 0.120, 95% CI: 0.075 to 0.164). Estimates of lnVR below zero (dashed grey line) indicate that there is less variability (SD) across gradient than uniform landscapes. Data points show the estimate of lnVR at each time point.

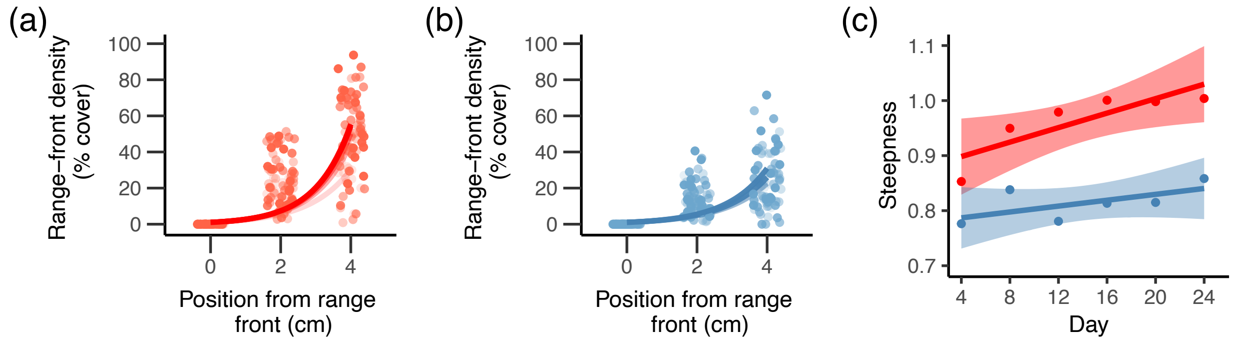


**Figure S4:** Steepness of the range front during range expansion across gradient (red) and uniform (blue). (a-b) Data points show estimates of population density (% cover) and lines show predictions from an exponential model of population density across space at the range front, with increasing colour intensity representing time (from day 4 to 24). (c) Lines and shading represent the slope and 95% CIs obtained from a linear model of steepness (i.e., the exponent) over time and by temperature treatment.

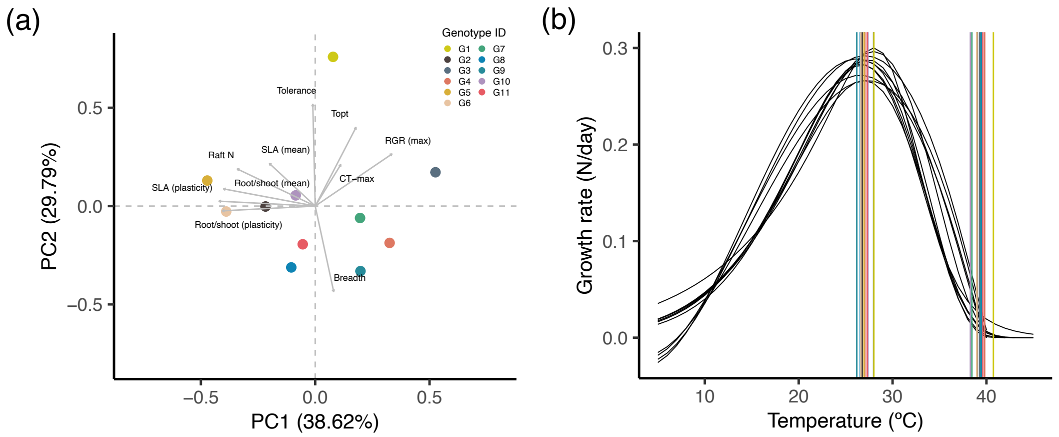


**Figure S5:** Trait variation among experimental genotypes. (a) PCA of trait space with the first two axes explaining 68.41% of the total variation. Coloured dots represent trait values for the 11 genotypes in trait space. Grey arrows represent the 10 measured traits: (1) thermal optimum ($T_{opt}$); (2) thermal performance breadth (Breadth); (3) thermal tolerance breadth (Tolerance); (4) critical thermal maximum ($CT_{max}$); (5) maximum growth rate ($RGR_{max}$); (6) mean SLA; (7) mean root/shoot ratio; (8) mean raft number; (9) plasticity in SLA ($SLA_{plasticity}$); and (10) plasticity in root/shoot ratio ($root/shoot_{plasticity}$). (b) Black curves show variation in thermal performance curves across the 11 genotypes. Vertical coloured lines represent the thermal optimum ($T_{opt}$) and critical thermal maximum ($CT_{max}$), with colours corresponding to the 11 genotypes in (a).

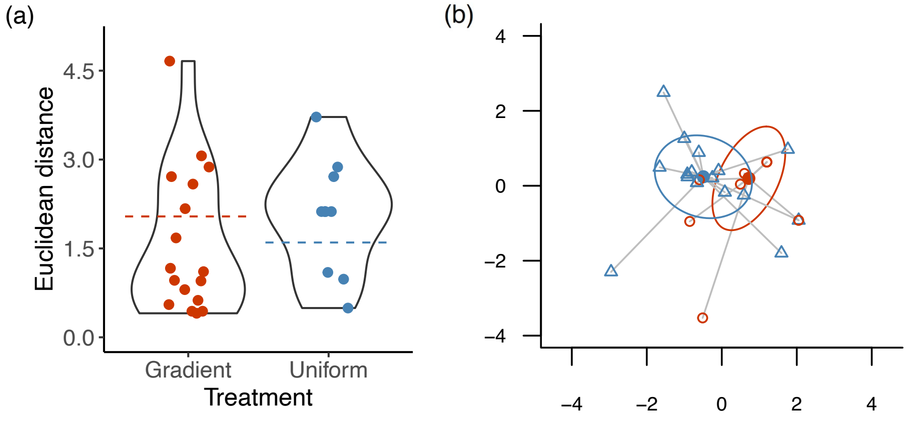


**Figure S6:** Trait change and composition in multidimensional trait space for range-front populations in gradient (red) and uniform (blue) landscapes. (a) Violin plots showing the extent of trait evolution. Jittered data points represent estimates of Euclidean distance between the final and initial trait value in each population for 10 traits. The dotted horizontal line shows the mean trait change observed. (b) Trait composition of range-front populations in gradient and uniform landscapes. The first two axes of a Principal Coordinates Analysis (PCoA) are shown, accounting for 67.2% of the total variation in trait composition. Filled circles are treatment centroids, while clear symbols are replicate populations in each treatment. Ellipses represent one standard deviation around treatment centroids, while the lines represent the distance between the centroid and each replicate.

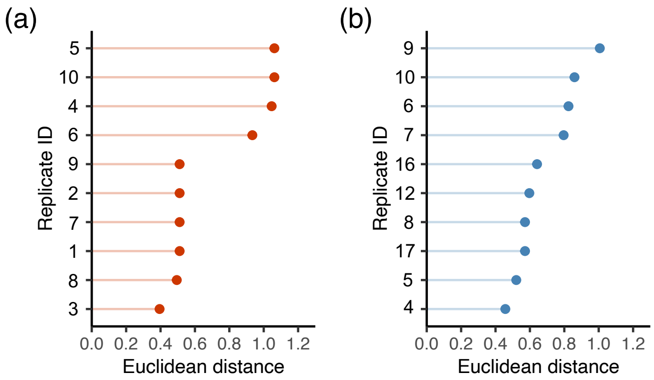


**Figure S7:** Euclidean distance between the mean genotype composition and the genotype composition of each replicate in (a) gradient and (b) uniform landscapes. Points represent the Euclidean distance for each replicate and lines are a visual aid for linking to each replicate ID. Data points are ordered from the most distant (top) to the least distant (bottom). Replicate IDs correspond to that of Figure S8 below.

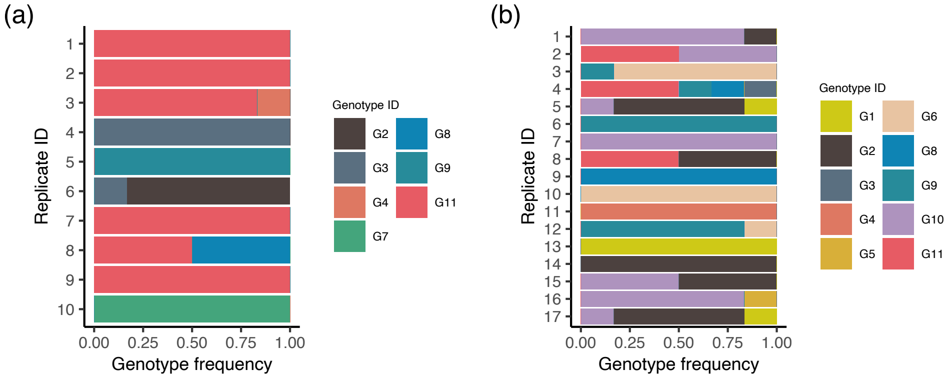


**Figure S8:** Within-treatment genotypic composition at the range front of populations expanding across (a) gradient and (b) uniform landscapes. Replicate IDs are placed from top to bottom in order of range expansion speed such that the top replicate ID travelled fastest. Colours represent unique genotype IDs.

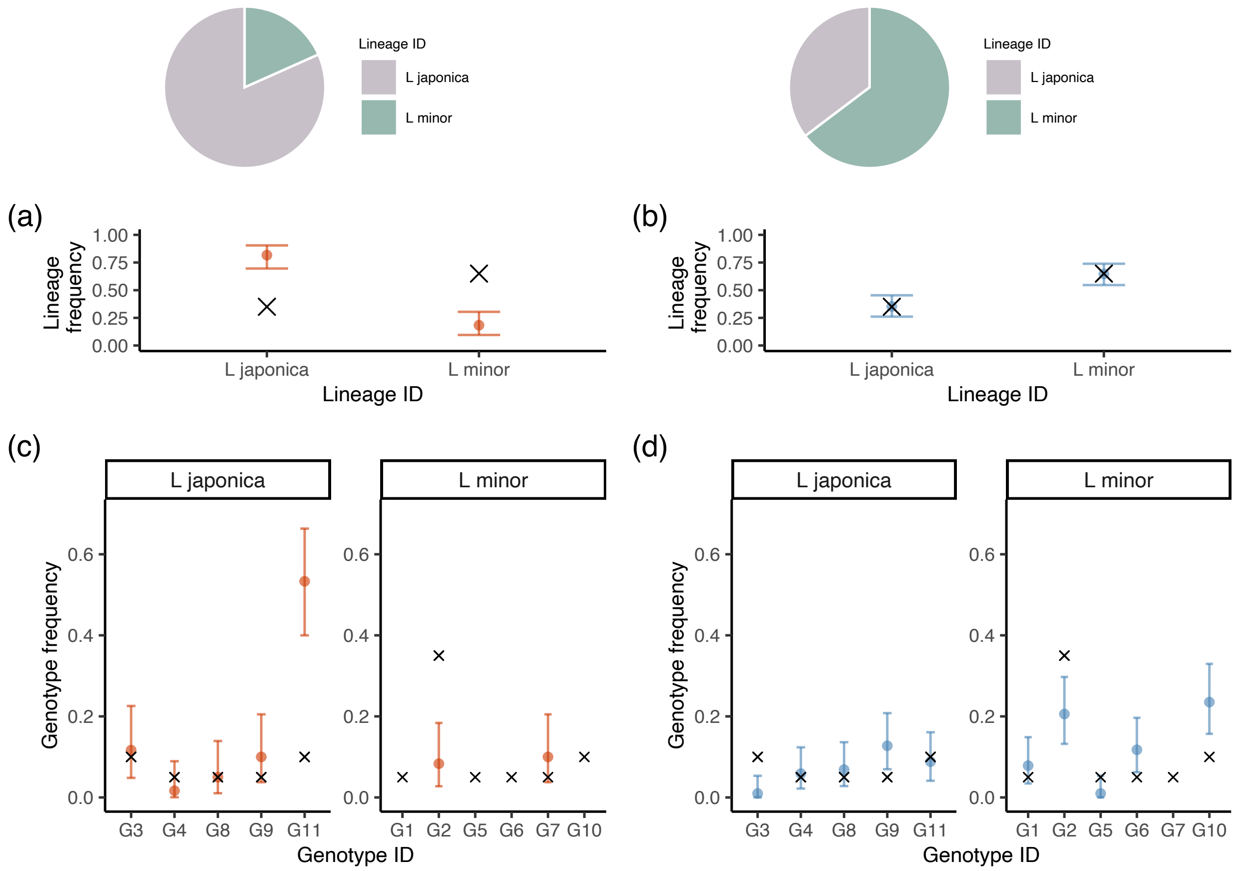


**Figure S9:** Composition of *L. minor* and *L. japonica* lineages for range-front populations in gradient and uniform landscapes as delimited by TBP markers. (a; b) Mean frequencies and 95% CI of *L. minor* and *L. japonica* observed across range-front populations in (a) gradient and (b) uniform landscapes. The initial "species" frequency in the founder population is denoted by 'x'. (c; d) Mean frequencies and 95% CI of genotypes observed across range-front populations in (c) gradient and (d) uniform landscapes. The initial genotype frequency in the founder population is denoted by 'x'. We note that the high frequency of *L. japonica* in gradient landscapes is driven by selection for genotype G11 rather than through an overall increase of all *L. japonica* genotypes.

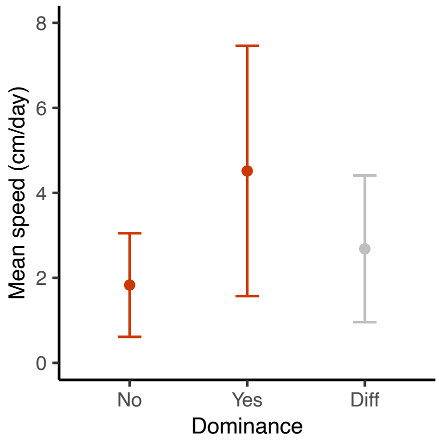


**Figure S10:** Mean speed (i.e., slope of distance over time) of gradient landscapes with and without G11 genotypes dominating at the range front. The points and error bars represent the mean and 95% CI, respectively. Dominance represents whether the genetic composition of range-front populations have a high proportion (>80%) of the G11 genotype or not (‘Yes’ or ‘No’, respectively). Diff = is the difference in mean speed between the two groups (slope = 2.684, 95% CI: 0.959 to 4.409 cm/day). Coefficients come from a linear model of distance travelled (cm) as the response variable and an interaction between time (day) and dominance (‘Yes’ or ‘No’) as fixed predictors ($N$ = 10 replicates).

**Table S1:** Sampling locations of *Lemna* accessions. Each row represents a uniquely sampled accession, with species identification based on TBP barcodes. Unique genotype IDs detected by microsatellite markers are shown. Genotype IDs in the brackets correspond to the genotype IDs across figures presented in the manuscript. Samples were collected between March and November in 2018 and 2019.

| Species | Genotype ID | Lat | Long | Location |
|---|---|---|---|---|
| *L. minor* | LM001 (G1) | 49.2639 | –123.2499 | Biodiversity Museum, Vancouver, BC |
| *L. minor* | LM002 (G2) | 49.2564 | –122.9648 | Burnaby Lake (Ditch), Burnaby, BC |
| *L. minor* | LM002 (G2) | 49.2565 | –122.9652 | Burnaby Lake (West), Burnaby, BC |
| *L. minor* | LM002 (G2) | 49.2175 | –123.1766 | Celtic Ave & Balaclava St, Vancouver, BC |
| *L. minor* | LM002 (G2) | 49.2184 | –123.1788 | Celtic Ave & Blenheim St, Vancouver, BC |
| *L. minor* | LM002 (G2) | 48.6975 | –122.4776 | Hemlock Trail, Bellingham, WA |
| *L. minor* | LM002 (G2) | 49.1738 | –123.1976 | Terra Nova Park (North), Richmond, BC |
| *L. minor* | LM002 (G2) | 49.2230 | –123.1762 | W53 Ave & Balaclava St, Vancouver, BC |
| *L. japonica* | LM003 (G3) | 49.2715 | –123.1107 | Hinge Park (North), Vancouver, BC |
| *L. japonica* | LM003 (G3) | 49.2706 | –123.1103 | Hinge Park (South), Vancouver, BC |
| *L. japonica* | LM004 (G4) | 49.2695 | –123.1358 | Granville Island, Vancouver, BC |
| *L. minor* | LM005 (G5) | 49.6952 | –124.5076 | Cranby Lake, Texada Island, BC |
| *L. minor* | LM006 (G6) | 48.7366 | –122.3820 | Geneva Pond, Bellingham, WA |
| *L. minor* | LM007 (G7) | 49.2658 | –123.2599 | Nitobe Memorial Garden, Vancouver, BC |
| *L. japonica* | LM008 (G8) | 49.2243 | –123.1759 | Southlands Heritage Farm, Vancouver, BC |
| *L. japonica* | LM009 (G9) | 49.2565 | –122.9647 | Burnaby Lake (North), Burnaby, BC |

| | | | | |
|---|---|---|---|---|
| *L. minor* | LM010 (G10) | 49.2522 | –122.9625 | Burnaby Lake (East), Burnaby, BC |
| *L. minor* | LM010 (10) | 49.1722 | –123.1984 | Terra Nova Park (South), Richmond, BC |
| *L. japonica* | LTU001 (G11) | 49.2382 | –122.9705 | Deer Lake (East), Vancouver, BC |
| *L. japonica* | LTU001 (G11) | 49.2383 | –122.9713 | Deer Lake (North), Vancouver, BC |

**Table S2:** Microsatellite markers used for genotyping accessions based on Hart et al. (2019). Allele size ranges (bp) are reported for accessions used in our experiment.

| Locus | Primer sequence (5'–3') | Repeat motif | Allele size range (bp) |
|---|---|---|---|
| R5C | F: TGATGCCAGTAGATCCGGC<br>R: ACGCCTGAACACGATTGATG | AGAT | 320 – 380 |
| R15A | F: GTGACAGCGTATCCTTGTGC<br>R: TCAGCGGCAAGATCATCAAG | ATC | 220 – 280 |
| R15B | F: TCGAGCTAATCAGTGGAGCC<br>R: TGAGTGCTCGGCTTGACTTTC | AG | 140 – 190 |
| R15C | F: TGTTCCCACCCACTTGAC<br>R: AAAGGAAGAGGGAGCAAGGG | AT | 370 – 390 |

**Table S3:** Primer sequence for barcoding accessions using $\beta$-tubulin genes.

| Locus | Primer sequence (5'–3') | Reference |
|---|---|---|
| *TUBB1* | F: CACYCCAAGCTGTAAGWTCC<br>R: GATCGCCGACTAYAAGAAATC | Braglia et al. 2021a |

**Table S4:** Model coefficients testing the difference in the mean speed (i.e., slope) of range expansion across gradient and uniform landscapes. Intercept represents maximum distance travelled for gradient landscapes (on Bench #1). Random effects include a random slope for each replicate ID. LCI and UCI correspond to lower and upper 95% confidence intervals, respectively. *Italicised* estimates have 95% confidence intervals that do not span zero.

| Fixed predictors | Estimate | LCI | UCI | *P* |
|---|---|---|---|---|
| Intercept | –13.737 | –28.535 | 1.956 | 0.073 |
| *Time (day)* | *2.499* | *1.083* | *3.916* | *0.002* |
| Treatment (uniform) | 1.414 | –12.294 | 14.888 | 0.845 |
| *Bench ID (#2)* | *17.633* | *1.368* | *33.898* | *0.049* |
| Bench ID (#3) | 3.306 | –17.276 | 23.888 | 0.736 |
| Bench ID (#4) | 4.537 | –15.573 | 24.646 | 0.601 |
| *Time * Treatment* | *2.584* | *0.610* | *4.558* | *0.016* |

**Table S5:** Model coefficients testing change in log coefficient of variation ratio (lnCVR) over time. Positive estimates of lnCVR indicate greater variability in distance travelled among gradient compared to uniform landscapes. Sensitivity models include sampling variance around lnCVR estimates in a meta-regression, with effect size IDs as a random effect. LCI and UCI correspond to lower and upper 95% confidence intervals, respectively. *Italicised* estimates have 95% confidence intervals that do not span zero.

| Model type | Fixed predictors | Estimate | LCI | UCI | *P* |
|---|---|---|---|---|---|
| Linear model | *Intercept* | *–1.220* | *–1.660* | *–0.779* | *0.002* |
| | *Time (day)* | *0.074* | *0.046* | *0.103* | *0.002* |
| Sensitivity model | *Intercept* | *–1.055* | *–1.490* | *–0.620* | *<0.001* |
| | *Time (day)* | *0.066* | *0.041* | *0.091* | *<0.001* |

**Table S6:** Model coefficients for tests on differences in range-front population density over time. Intercept represents percent cover (%) for gradient landscapes on day 0. LCI and UCI correspond to lower and upper 95% confidence intervals, respectively. *Italicised* estimates have 95% confidence intervals that do not span zero.

| Model type | Fixed predictors | Estimate | LCI | UCI | *P* |
|---|---|---|---|---|---|
| Linear model | *Intercept* | *25.771* | *17.319* | *34.224* | *<0.001* |
| | *Time (day)* | *2.311* | *1.768* | *2.853* | *<0.001* |
| | Treatment (uniform) | –3.011 | –14.964 | 8.943 | 0.577 |
| | *Time * Treatment* | *–1.460* | *–2.228* | *–0.693* | *0.002* |
| Sensitivity model | *Intercept* | *24.995* | *17.824* | *32.167* | *<0.001* |
| | *Time (day)* | *2.359* | *1.883* | *2.835* | *<0.001* |
| | Treatment (uniform) | –2.719 | –12.854 | 7.417 | 0.599 |
| | *Time * Treatment* | *–1.485* | *–2.169* | *–0.801* | *<0.001* |

**Table S7:** Model coefficients for tests on differences in steepness (i.e., rate of change in population density) at the range front. Intercept represents the exponent obtained from an exponential model of population density over space for gradient landscapes on day 0. LCI and UCI correspond to lower and upper 95% confidence intervals, respectively. *Italicised* estimates have 95% confidence intervals that do not span zero.

| Model type | Fixed predictors | Estimate | LCI | UCI | *P* |
|---|---|---|---|---|---|
| Linear model | *Intercept* | *0.872* | *0.805* | *0.939* | *<0.001* |
| | *Time (day)* | *0.007* | *0.002* | *0.011* | *0.008* |
| | *Treatment (uniform)* | *–0.096* | *–0.191* | *–0.001* | *0.049* |
| | Time * Treatment | –0.004 | –0.010 | 0.002 | 0.177 |
| Sensitivity model | *Intercept* | *0.872* | *0.814* | *0.930* | *<0.001* |
| | *Time (day)* | *0.007* | *0.003* | *0.010* | *<0.001* |
| | *Treatment (uniform)* | *–0.096* | *–0.176* | *–0.015* | *0.020* |
| | Time * Treatment | –0.004 | –0.009 | 0.002 | 0.162 |

**Table S8:** Model coefficients for tests on the extent of change in genotype or trait frequencies, as quantified by Euclidean distance between evolved and founder populations. Linear models were fitted separately for genotype and trait frequencies. Intercept represents the Euclidean distance for gradient landscapes on Bench ID #1. LCI and UCI correspond to lower and upper 95% confidence intervals, respectively. *Italicised* estimates have 95% confidence intervals that do not span zero.

| Response | Fixed predictors | Estimate | LCI | UCI | *P* |
|---|---|---:|---:|---:|---:|
| Genotype | *Intercept* | *0.820* | *0.615* | *1.026* | *<0.001* |
| | Treatment (uniform) | –0.121 | –0.295 | 0.053 | 0.163 |
| | *Bench ID (#2)* | 0.295 | –0.004 | 0.594 | 0.053 |
| | Bench ID (#3) | 0.020 | –0.205 | 0.244 | 0.857 |
| | Bench ID (#4) | 0.131 | –0.089 | 0.351 | 0.229 |
| Trait | *Intercept* | *1.834* | *0.723* | *2.946* | *0.002* |
| | Treatment (uniform) | –0.452 | –1.393 | 0.490 | 0.330 |
| | Bench ID (#2) | 1.296 | –0.322 | 2.913 | 0.111 |
| | Bench ID (#3) | 0.160 | –1.056 | 1.375 | 0.788 |
| | Bench ID (#4) | 0.061 | –1.130 | 1.251 | 0.917 |

**Table S9:** Model coefficients from PERMANOVA of genotype or trait composition. Bench ID was included as a fixed covariate. *SS* = sum of squares. *Italicised* estimates have $P < 0.05$.

| Response | Source | *Df* | *SS* | $R^2$ | *F* | *P* |
|---|---|---|---:|---:|---:|---:|
| Genotype | *Treatment* | *1* | *70.9* | *0.111* | *3.269* | *0.002* |
| | Bench ID | 3 | 92.8 | 0.145 | 1.425 | 0.089 |
| | Residual | 22 | 477.3 | 0.745 | | |
| | Total | 26 | 641.0 | | | |
| Trait | Treatment | 1 | 9.0 | 0.080 | 2.119 | 0.066 |
| | Bench ID | 3 | 10.6 | 0.093 | 0.828 | 0.635 |
| | Residual | 22 | 93.9 | 0.827 | | |
| | Total | 26 | 113.6 | | | |

**Table S10:** Final trait values in uniform and gradient landscapes. Trait values are scaled, centred and weighted by the final genotype frequency of each population. LCI and UCI correspond to lower and upper 95% confidence intervals, respectively. *Italicised* estimates have 95% confidence intervals that do not span the initial trait value of the founder population.

| Treatment | Trait | Mean | LCI | UCI |
|---|---|---|---|---|
| Uniform | Thermal breadth | –0.015 | –0.357 | 0.327 |
| | CT-max | –0.054 | –0.413 | 0.305 |
| | RGR-max | 0.018 | –0.244 | 0.281 |
| | Raft number | 0.096 | –0.256 | 0.448 |
| | Root/shoot | 0.039 | –0.089 | 0.166 |
| | Root/shoot plasticity | 0.213 | –0.121 | 0.546 |
| | SLA | –0.056 | –0.295 | 0.183 |
| | SLA plasticity | 0.209 | –0.027 | 0.445 |
| | Tolerance | –0.033 | –0.402 | 0.335 |
| | Thermal optimum | –0.130 | –0.461 | 0.202 |
| Gradient | *Thermal breadth* | *0.602* | *0.119* | *1.085* |
| | CT-max | 0.129 | –0.203 | 0.461 |
| | RGR-max | –0.402 | –0.988 | 0.184 |
| | Raft number | –0.328 | –0.663 | 0.007 |
| | Root/shoot | –0.243 | –0.620 | 0.133 |
| | Root/shoot plasticity | –0.294 | –0.614 | 0.027 |
| | SLA | –0.250 | –0.543 | 0.043 |
| | SLA plasticity | –0.070 | –0.564 | 0.425 |
| | *Tolerance* | *–0.407* | *–0.646* | *–0.169* |
| | Thermal optimum | 0.143 | –0.342 | 0.628 |

**Table S11:** Model coefficients from *t*-tests on differences in trait means between gradient and uniform landscapes. *Italicised* estimates have $P < 0.05$.

| Trait | *t* | *Df* | *P* |
|---|---|---|---|
| Thermal breadth | 2.043 | 17.782 | 0.056 |
| CT-max | 0.737 | 23.937 | 0.469 |
| RGR-max | –1.711 | 23.642 | 0.100 |
| Raft number | –1.282 | 12.687 | 0.223 |
| Root/shoot | –1.391 | 11.101 | 0.192 |
| *Root/shoot plasticity* | *–2.147* | *23.527* | *0.042* |
| SLA | –1.007 | 19.977 | 0.326 |
| SLA plasticity | –0.995 | 13.189 | 0.338 |
| Tolerance | –1.671 | 24.549 | 0.107 |
| Thermal optimum | 0.910 | 17.257 | 0.375 |

**Table S12:** Model coefficients for change in genotype frequencies at the range core and at the range edge from the founder population. Amount of change in genotype frequency was quantified as the Euclidean distance between evolved and founder populations. Linear models were fitted separately for gradient and uniform landscapes. Intercept represents the Euclidean distance for range core populations on Bench ID #1. LCI and UCI correspond to lower and upper 95% confidence intervals, respectively. *Italicised* estimates have 95% confidence intervals that do not span zero.

| Treatment | Fixed predictors | Estimate | LCI | UCI | *P* |
|---|---|---|---|---|---|
| Gradient | Intercept | 0.154 | –0.033 | 0.342 | 0.091 |
| | *Site (edge)* | *0.511* | *0.358* | *0.664* | *<0.001* |
| | Bench ID (#3) | 0.294 | 0.096 | 0.492 | 0.011 |
| | Bench ID (#4) | 0.307 | 0.065 | 0.549 | 0.021 |
| Uniform | *Intercept* | *0.509* | *0.194* | *0.823* | *0.007* |
| | Site (edge) | 0.213 | 0.123 | 0.549 | 0.171 |
| | Bench ID (#3) | 0.008 | –0.367 | 0.384 | 0.959 |
| | Bench ID (#4) | 0.107 | –0.353 | 0.567 | 0.589 |
